# Asymmetric condensin loop extrusion is regulated by RPA-coated single-stranded DNA in quiescent cells

**DOI:** 10.64898/2026.07.10.737861

**Authors:** Ban Al-Kurdi, Jason A. Hernandez, Annabel H. Lewis, Lauren M. Snyder, Steven M. Markus, Sarah G. Swygert

## Abstract

SMC complexes influence virtually all DNA-dependent processes by organizing the genome via the process of loop extrusion. Although transcription has been implicated in regulating SMC complex function, the underlying mechanisms remain unclear. Further, the directionality of loop extrusion observed in biochemical experiments has been difficult to reconcile with the chromatin condensation observed in cells. Here, we use a quiescent yeast model to uncover the relationship between condensin loop extrusion and transcription. Condensin gradually relocates to transcribed gene promoters during quiescence entry, allowing us to dissect condensin targeting mechanisms temporally. Through targeted degradation experiments, we discover that topological stress generated by transcription leads to single-stranded DNA accumulation at promoters, and that these RPA-bound regions are loading sites and extrusion barriers for condensin. We further use a condensin mutant to determine that condensin extrudes loops asymmetrically in cells. We propose that antagonism by RPA universally regulates SMC complex function.

## INTRODUCTION

The structural maintenance of chromosome (**SMC**) complexes, cohesin, condensin, and SMC5/6, are highly conserved, essential protein complexes implicated in regulating nearly every DNA-dependent process.^1–5^ SMC complexes organize the three-dimensional (**3D**) structure of the genome by creating loops of chromatin that condense chromosomes during mitosis,^6–11^ dictate promoter/enhancer interactions during interphase,^12–14^ and facilitate DNA repair.^15–19^ SMC complexes are believed to generate chromatin loops through loop extrusion,^20–25^ during which the complex loads onto DNA and reels chromatin in an ATP-dependent manner to elongate the loop.^1,2,4^ Loops are extended until reaching a barrier, leading to transient stabilization.^1,4,5^ Stabilized loops are referred to as chromatin loop domains, regions of associating chromatin flanked by insulating boundaries where the SMC complex is bound. Although loop extrusion has been visualized in biochemical single-molecule studies of all three SMC complexes,^22–25^ the extrusion mechanisms observed *in vitro* have been difficult to reconcile with the extent of chromatin compaction in cells.^26^

While extrusion barriers have been identified for SMC complexes in certain situations, such as CTCF serving as a barrier for cohesin in interphase mammalian cells,^27–29^ it is generally unclear why SMC complexes stall when encountering some obstacles while bypassing others. Of these obstacles, transcription is particularly confounding. The interplay between transcription and SMC-mediated chromatin folding is conserved, as domain boundaries overlap with active transcription across organisms.^30–36^ In yeast and mammalian cells, transcription either pushes cohesin^30,37–39^ or serves as a moving extrusion barrier,^32,40^ leading to cohesin accumulation at the 3’ ends of convergently oriented genes. Cohesin has been proposed to load at active genes,^41–43^ however a more recent study suggests that it must load genome-wide to compact the genome.^44^ The moving barrier model has also been proposed to regulate bacterial and fission yeast condensin.^32,40,45–47^ However, budding yeast condensin can bypass RNA polymerase in biochemical experiments.^48^ Consequently, although transcription is heavily implicated in regulating SMC complex dynamics, the underlying mechanisms remain unclear.

To address these questions, we utilized quiescent *Saccharomyces cerevisiae* as model to investigate condensin-mediated loop extrusion. Quiescence is a conserved state in which cells reversibly exit the cell cycle and undergo massive transcriptional shut down and metabolic reprograming.^49–55^ Quiescence is essential for maintaining stem cell repertoires and is implicated in chemotherapeutic evasion and cancer recurrence.^49,56–58^ Budding yeast enter quiescence in response to nutrient deprivation.^51,59^ As cultured cells exhaust glucose in the media, young cells enter quiescence gradually over several days, and these cells can be purified from non-quiescent cells by separation on density gradients.^51,59^ We have previously shown that between cycling and quiescence, condensin relocalizes from tRNA genes, centromeres, and the rDNA to bind coding gene promoters, forming chromatin loop domains responsible for compacting the genome and repressing approximately 20% of all genes.^60^ This condensin-dependent chromatin condensation is conserved in quiescent mammalian fibroblasts and lymphocytes.^60,61^

Here, we show that condensin accumulates at transcribed gene promoters during quiescence entry over several days, allowing for temporal dissection of condensin mechanisms. This condensin relocalization is regulated by transcription-dependent topological stress that generates regions of Replication Protein A (**RPA**)-coated single-stranded DNA (**ssDNA**) at active promoters during quiescence. These regions serve both as loading sites and extrusion barriers for condensin, which loads at RPA-coated promoter regions and extrudes loops asymmetrically until reaching a subsequent RPA binding site. Collectively, these results reveal the mechanisms by which transcription regulates condensin loop extrusion and confirm predictions of condensin extrusion mechanics proposed by modeling and biochemical single-molecule studies in cells.

## RESULTS

### Condensin gradually accumulates at coding gene promoters during quiescence entry

We have previously shown that during quiescence entry, condensin relocates to coding gene promoters, where it stabilizes large chromosomal interaction domains (**L-CID**s) as repressive loops.^60^ This process coincides with dramatic condensation of the genome. By convention, *S. cerevisiae* cultures are inoculated for quiescence on Day 0 and purified on Day 7, though cells enter stationary phase at the diauxic shift by Day 1, giving cells nearly a week after cycling to enter quiescence.^51^ We wondered if condensin relocalization and chromatin condensation coincide during this time. To test this, we completed confocal microscopy and chromatin volume measurements of quiescent cells purified from Day 3 to Day 7 using strains in which one copy of H2B is tagged with GFP (**Fig. 1A**). In this and all subsequent experiments, samples were performed in biological replicate in isogenic strains derived from independent parental strains. Our analysis revealed a gradual reduction in mean chromatin volume from Day 3 to Day 7 at the population level (**Fig. 1B**), with maximal condensation achieved by Day 6.

**Figure 1.**
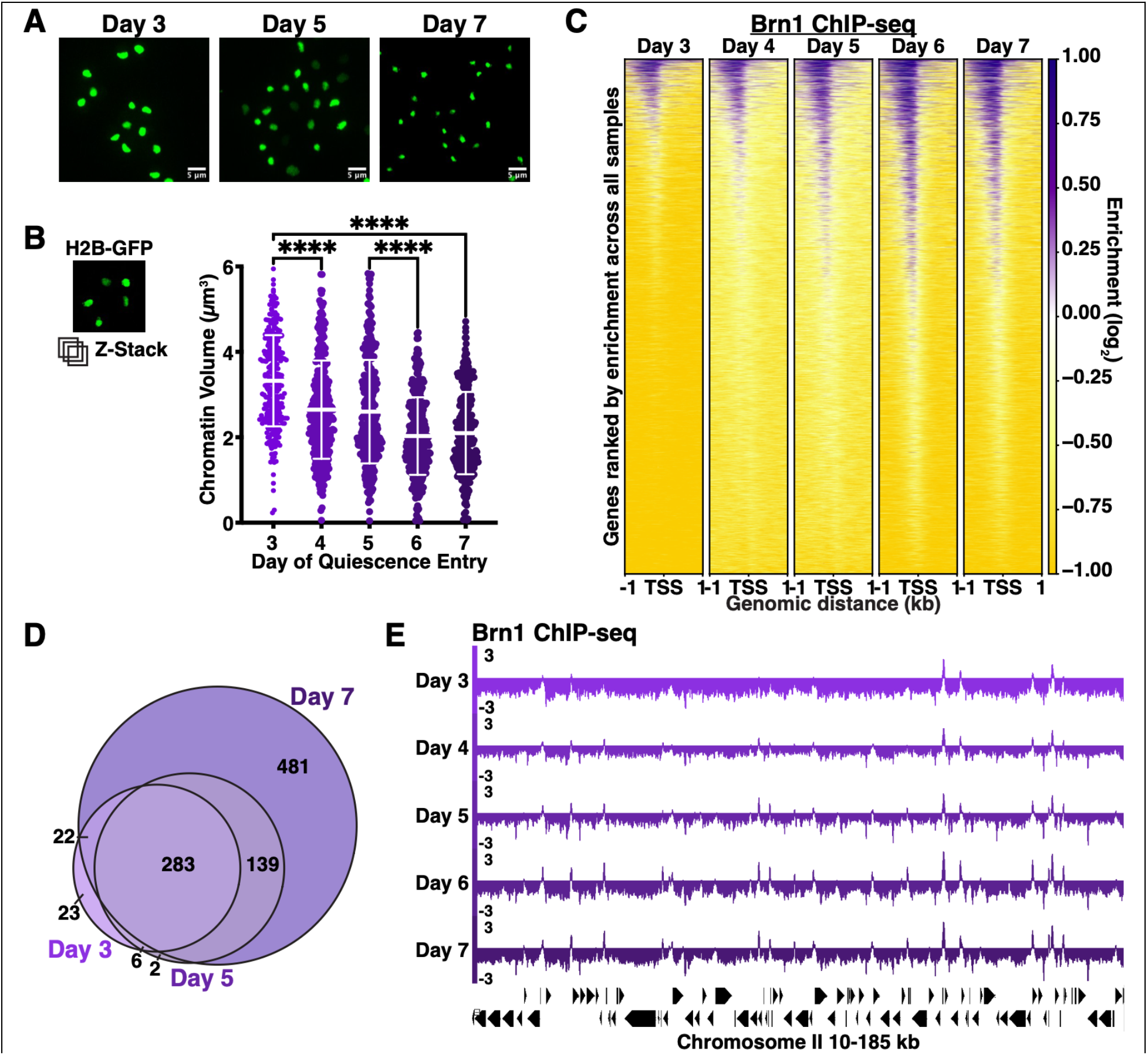
Condensin gradually accumulates at promoters during quiescence entry. **(A)** Representative confocal microscopy images of H2B-GFP quiescent cells purified on the indicated day. Scale bar, 5 µm. (**B**) Chromatin volume quantification. Bars represent mean +/- SD. Each time point consists of at least 150 cells from each of two biological replicates. Significance measured using one-way ANOVA. (**C**) ChIP-seq heatmaps of Brn1- FLAG in quiescent cells purified on the indicated day. For all ChIP-seq experiments and analyses, data from two biological replicates were merged. Heatmaps show all annotated RNAPII transcripts in the same order unless otherwise indicated. (**D**) Overlap of ChIP-seq peaks. Peaks were filtered using a 1.5-fold enrichment threshold and direct overlaps were determined. (**E**) Representative genome browser view of ChIP-seq data.

We next examined condensin binding during this same time period by completing ChIP-seq of strains in which the Brn1 subunit of condensin is tagged with FLAG. In all ChIP-seq experiments, data from biological replicates were confirmed to agree and merged to increase signal to noise. This time course reveals that condensin gradually accumulates at coding gene promoters during quiescence entry, with maximal binding reached by Day 6, consistent with the timeframe of chromatin condensation (**Fig. 1C**). To quantify changes in occupancy and to compare condensin binding sites throughout the time course, we called Brn1 ChIP-seq peaks and examined their overlap (**Fig. 1D**).^62,63^ We found that the number of peaks increases between Day 3 and Day 7, and that most early peaks overlap with peaks at Day 7. Consistently, examination at the genome browser level demonstrates that condensin peaks appear at varying times and then increase in magnitude over several days (**Fig. 1E**). Overall, these results suggest that chromatin condensation during quiescence entry coincides with the gradual accumulation of condensin at promoters, and that quiescence entry can serve as a model system to dissect condensin dynamics temporally without the confounding influence of cell cycle progression.

### Transcription targets condensin to active gene promoters during quiescence entry

Although most genes are silenced during quiescence, we have previously found that condensin is enriched at the promoters of the subset of genes that are active.^55,62^ We hypothesized that transcription might play a role in condensin localization to those sites. To test this, we performed ChIP-seq of Rpb1, the largest RNA Polymerase II (**RNAPII**) subunit, during quiescence entry (**Fig. S1A**). We observed that, although transcription is dramatically reduced in quiescent cells compared to cycling cells,^53,60,64^ transcription changes dynamically during Days 3-7, with some regions losing Rpb1 while others retain or even gain Rpb1 as cells progress further into quiescence. Next, we compared changes in Rpb1 occupancy to Brn1 occupancy during this time period (**Fig. 2A-B**). We found that condensin binds transcribed promoters throughout the time course, with Rpb1 peaks preceding most Brn1 peaks. These results suggest that condensin preferentially binds transcribed genes, with transcription occurring prior to condensin accumulation at promoters.

**Figure 2.**
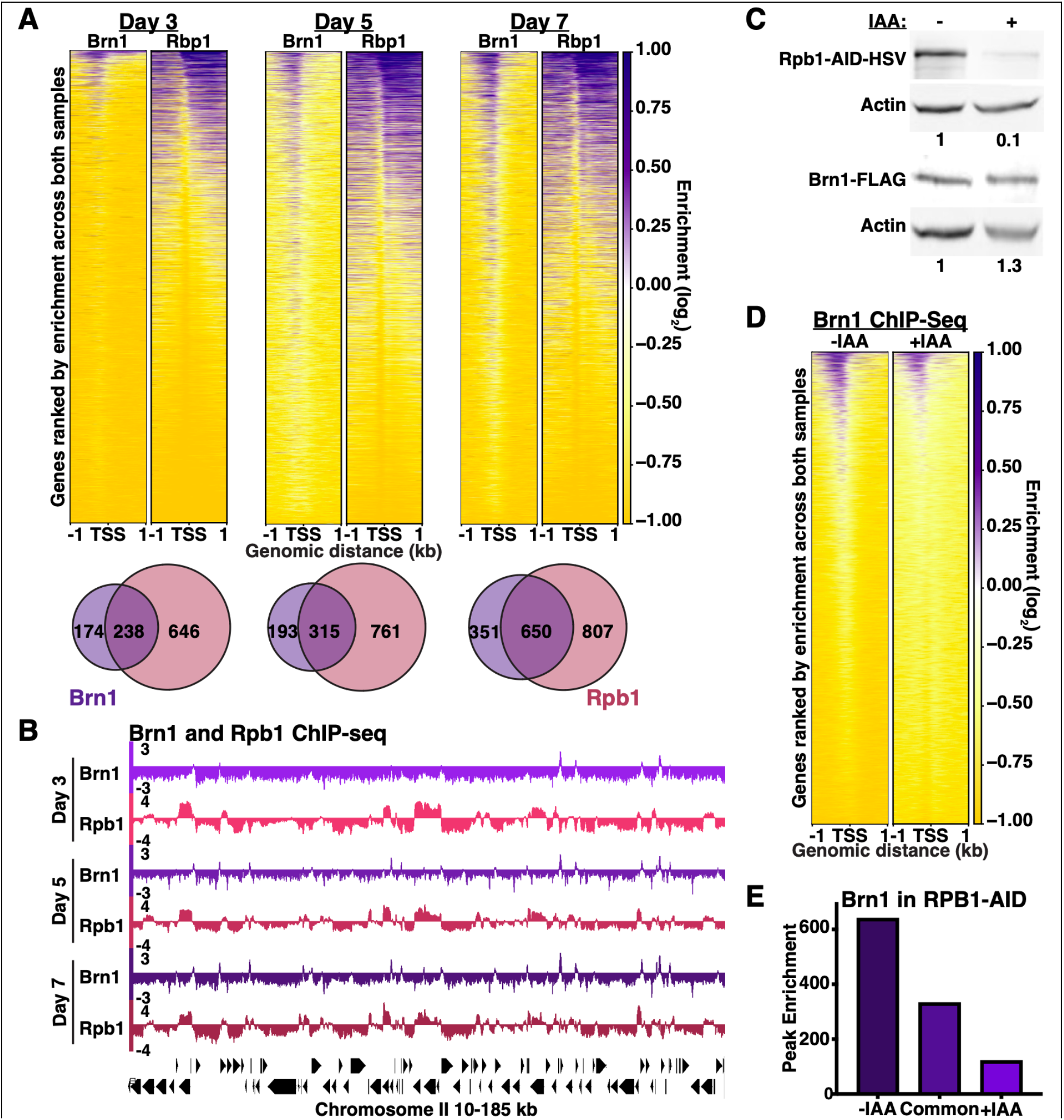
Transcription targets condensin to active gene promoters. **(A)** Top: ChIP-seq heatmaps of Brn1- FLAG and Rpb1 in quiescent cells purified on the indicated day. Bottom: Overlap of ChIP-seq peaks filtered using a 1.5-fold enrichment threshold. Peaks within 1 kb were determined to overlap. (**B**) Representative genome browser view of ChIP-seq data. (**C**) Representative Western blot in RPB1-AID-HSV Day 7 cells +/- IAA. Bands were normalized to actin and then to -IAA. (**D**) ChIP-seq heatmaps of Brn1-FLAG in RPB1-AID-HSV Day 7 cells +/- IAA. (**E**) Differential peak analysis of Brn1-FLAG in RPB1-AID-HSV Day 7 cells +/- IAA.

To test if transcription targets condensin during quiescence, we constructed auxin-inducible degron (**AID**) strains in which Rpb1 undergoes targeted degradation upon supplementation with indole-3-acetic acid (**IAA**).^65^ Addition of IAA during quiescence entry allows for efficient depletion of Rpb1 without affecting Brn1 protein levels when measured at Day 7 (**Fig. 2C**). To test the effect of Rpb1 depletion on condensin localization, we repeated Brn1 ChIP-seq at Day 7 in the *RPB1*-AID strains. In the presence of IAA, Brn1 occupancy at promoters is strongly reduced (**Fig. 2D**). As the peaks remaining in the presence of IAA overlap with peaks in the absence of IAA, we calculated differential peak enrichment, which determines differences in both peak location and magnitude across conditions so that peaks of similar enrichment at the same locations are considered to be common (**Fig. 2E**).^62^ This analysis confirms a strong reduction in Brn1 peak enrichment in the absence of Rpb1. These results suggest that during quiescence entry, transcription targets condensin to coding gene promoters.

### RPA-coated ssDNA accumulates at coding gene promoters during quiescence

We next sought to understand the mechanism by which transcription directs condensin during quiescence. In budding yeast, transcription has been proposed to influence cohesin localization by acting as a moving barrier to loop extrusion, resulting in the concentration of cohesin at the 3’ end of genes.^32,40^ In contrast, we find condensin at the 5’ end of genes, suggesting that transcription targets condensin through a distinct mechanism. According to the twin domain model, elongation generates positive supercoils ahead of the transcription bubble and negative supercoils behind it.^66,67^ As previous studies have shown that condensin preferentially binds supercoiled DNA,^68^ we hypothesized that transcription-induced topological stress may contribute to condensin relocalization in quiescence.

In cycling cells, transcription-generated supercoiling is resolved by Topoisomerases I and II (Top1 and 2 in *S. cerevisiae*).^66,69–71^ However, to our knowledge, Top1 and 2 activity had not previously been examined in quiescent budding yeast. To determine if Top1 and 2 are expressed during quiescence, we performed Western blots of Top1 and Top2 proteins tagged with FLAG in both cycling and quiescent cells (**Fig. 3A**). As we are unaware of a protein whose levels do not change been cycling and quiescence, we used both actin and H2B as loading controls and loaded the same amount of protein in each lane. We found that both Top1 and 2 show a remarkable reduction in protein levels in quiescent cells, suggesting that the ability to resolve transcription-induced supercoiling is limited in quiescence. Transcriptional elongation through highly compacted chromatin fibers may further induce topological stress during quiescence.^64^

**Figure 3.**
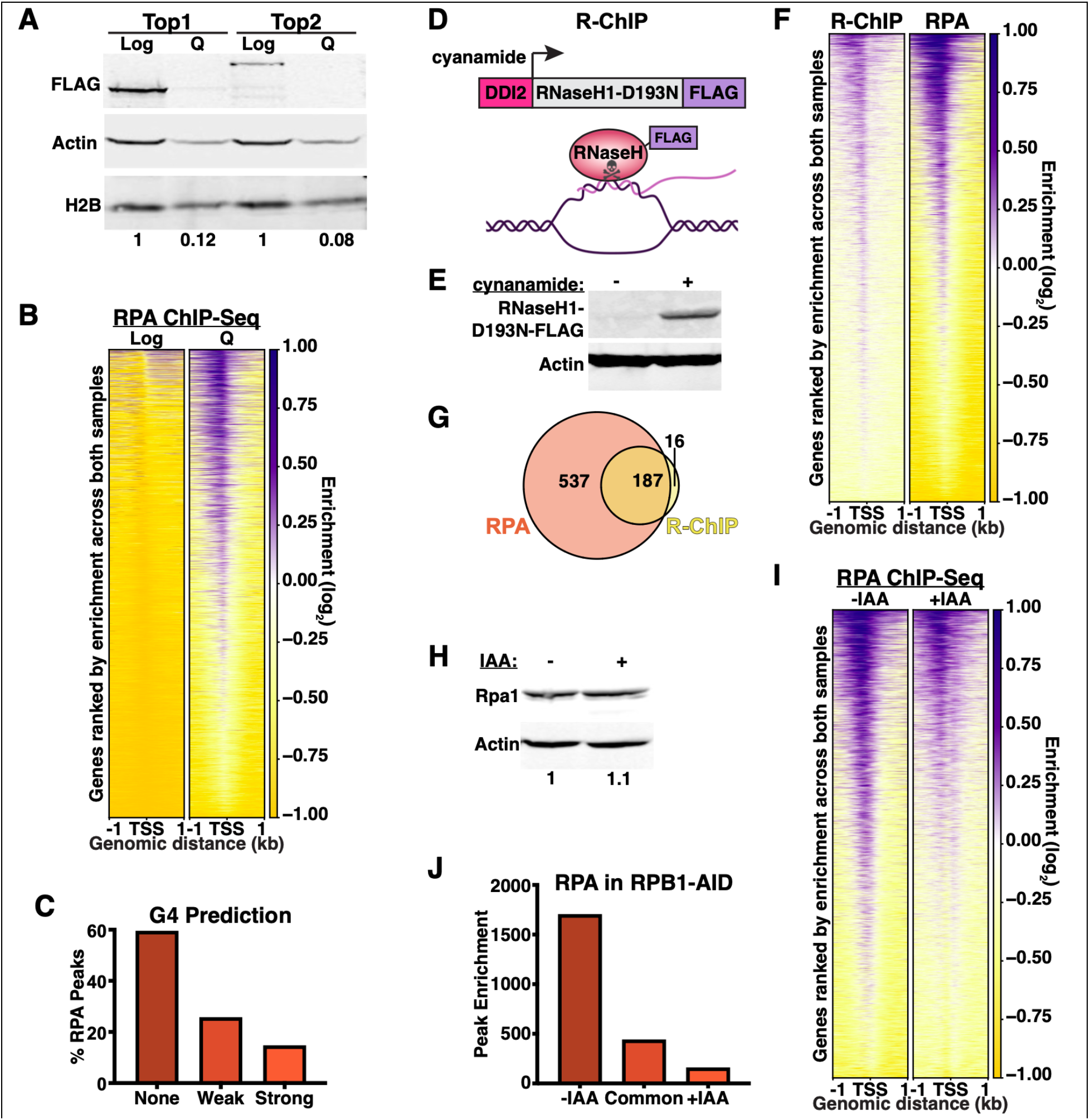
Topological stress accumulates at promoters during quiescence. **(A)** Representative Western blot of Top1-FLAG and Top2-FLAG in cycling (**log**) and Day 7 quiescent (**Q**) cells. Equal amounts of protein were added per lane. Bands were normalized to actin and then to log. (**B**) ChIP-seq heatmaps of RPA in log and Day 7 quiescent cells. (**C**) G4 predictions of RPA-bound sequences. “None” indicates a G4Hunter score < 1.2, “weak” indicates a score between 1.2 and 1.5, and “strong” indicates a score > 1.5. (**D**) Cartoon depicting the R-ChIP method. (**E**) Representative Western blot from Day 7 RNaseH1-D193N-FLAG cells +/- cyanamide. (**F**) ChIP-seq heatmaps of R-loops and RPA at Day 7. (**G**) Overlap of R-ChIP and RPA ChIP-seq peaks filtered using a 1.5-fold enrichment threshold. Peaks within 1 kb were determined to overlap. (**H**) Representative Western blot in *RPB1*-AID-HSV Day 7 cells +/- IAA. Bands were normalized to actin and then to -IAA. (**I**) ChIP-seq heatmaps of RPA in *RPB1*-AID-HSV Day 7 cells +/- IAA. (**J**) Differential peak analysis of RPA in *RPB1*-AID-HSV Day 7 cells +/- IAA.

Unresolved torsional stress results in the generation of underwound DNA near the 5’ end of genes. In cells, unwound ssDNA is coated with RPA, which can be used as a proxy to detect ssDNA^72^. RPA is a heterotrimeric protein complex consisting of Rpa1, Rpa2, and Rpa3.^72^ To determine if ssDNA forms at promoters during quiescence, we performed RPA ChIP-seq using a commercial polyclonal antibody against the complex (**Fig. 3B**). We found a striking accumulation of RPA at promoters in quiescent cells as compared to cycling, suggesting that topological stress accumulates at promoters during quiescence.

ssDNA can be stabilized by the formation of G4-quadruplexes (**G4**s).^73^ To test if G4 formation contributes to ssDNA accumulation during quiescence, we examined the G4-forming potential of RPA-bound sequences (**Fig. 3C**).^74^ We found that ∼40% of RPA peaks contain sequences with the potential to form G4s. The accumulation of negative supercoiling behind elongating RNA polymerases can also promote RNA:DNA hybrid (**R-loop**) formation, which can stabilize ssDNA on the non-hybridized side and absorb some torsional stress.^75,76^ To determine if R-loops are present at RPA sites, we performed R-ChIP.^77,78^ R-ChIP tracks the occupancy of a catalytically dead RNaseH1 protein that can bind but not resolve R-loops to map the locations of R-loops in cells (**Fig. 3D**). To allow for induction of the R-ChIP target during quiescence, we constructed strains with a copy of *RNH1-D193N*-FLAG under the control of the cyanamide-inducible *DDI2* promoter integrated at the *URA3* locus.^79^ Treating cells with cyanamide during quiescence entry efficiently induced RNaseH1-D193N expression (**Fig. 3E**). R-ChIP was then completed via ChIP-seq against FLAG, and the overlap between R-ChIP and RPA peaks was calculated (**Fig. 3F-G**). We found that the vast majority of R-loops overlap with RPA sites, as expected, and that ∼35% of RPA sites overlap with R-loops. We conclude that ssDNA accumulates at promoters during quiescence due to a non-mutually exclusive combination of DNA unwinding as a result of negative supercoiling, G4 formation, and R-loops.

To validate that RPA occupancy at promoters is due to transcription-induced topological stress, we examined RPA in Day 7 *RPB1*-AID cells in the presence and absence of IAA. We found that Rpa1 protein levels are not affected by Rpb1 depletion (**Fig. 3H**), however RPA occupancy at promoters is strongly reduced (**Fig. 3I-J**). Collectively, these results suggest that RPA accumulates at promoters during quiescence due to transcription-induced topological stress resulting from downregulation of Top1 and Top2.

### RPA-coated ssDNA strongly overlaps with condensin sites during quiescence entry

We hypothesized that transcription may be directing condensin localization by generating topological stress at promoters, as condensin has been shown to target RPA-bound ssDNA both in biochemical experiments and in cells.^80,81^ To test this, we performed ChIP-seq of RPA at Days 3, 5, and 7 during quiescence entry and compared the data to the Brn1 ChIP-seq data at these time points (**Fig. 4A**). We found that RPA gradually accumulates at the same promoters at approximately the same time as condensin, with nearly all RPA peaks overlapping with Brn1 at Day 7. Although the peak calling reveals extensive colocalization, we felt it was underestimating the overlap between Brn1 and RPA at promoters that we could see by examination at the genome browser level (**Fig. 4B**), possibly due to using a stringent enrichment threshold to call peaks. To assess overlap independently of peak calling, we sorted heatmaps solely based on Brn1 or RPA occupancy rather than shared occupancy (**Fig. S2A**). Regardless of the sorting method, heatmaps appear nearly identical, suggesting nearly full overlap at RNAPII genes. In contrast, comparing Brn1 and Rpb1 colocalization by differential heatmap sorting reveals a somewhat weaker correlation than between Brn1 and RPA (**Fig. S2B**). We also performed k-means clustering of Brn1, RPA, and Rpb1 ChIP-seq signals at Day 7 and found that although most clusters containing Brn1 also have RPA and Rpb1, genes within Cluster 2 demonstrated substantially less Rpb1 occupancy as compared to Brn1 and RPA (**Fig. 4C**). We speculate that Cluster 2 genes may have been expressed prior to our time course, with torsional stress persisting at these sites at Day 7. These results suggest that RPA is a better predictor of Brn1 occupancy than Rpb1, consistent with our hypothesis that transcription targets condensin through generation of RPA-coated ssDNA.

**Figure 4.**
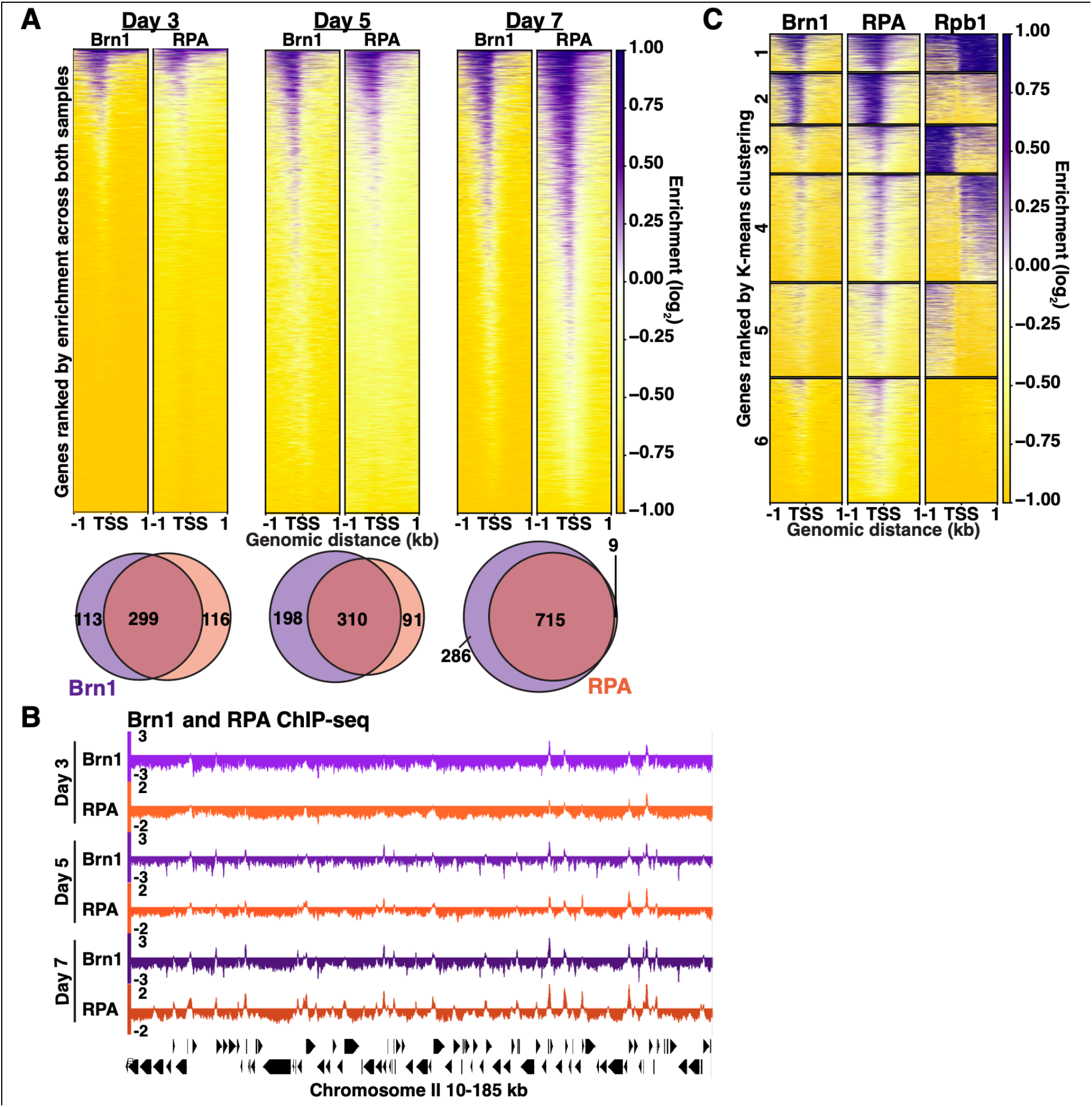
RPA-coated ssDNA overlaps spatially and temporally with condensin sites. **(A)** Top: ChIP-seq heatmaps of Brn1-FLAG and RPA in quiescent cells purified on the indicated day. Bottom: Overlap of ChIP-seq peaks filtered using a 1.5-fold enrichment threshold. Peaks within 1 kb were determined to overlap. (**B**) Representative genome browser view of ChIP-seq data. (**C**) K-means clustering of Brn1-FLAG, RPA, and Rpb1 ChIP-seq data from Day 7 cells.

### RPA, not transcription, is required for condensin targeting to promoters

To test if RPA is required for condensin targeting, we constructed AID strains of the largest RPA subunit, Rpa1, encoded by the *RFA1* gene. Addition of IAA during quiescence entry results in Rpa1 depletion but does not affect Brn1 protein levels as measured on Day 7 (**Fig. 5A**). We next completed ChIP-seq of Brn1 in the *RFA1*-AID strains purified on Day 7.

**Figure 5:**
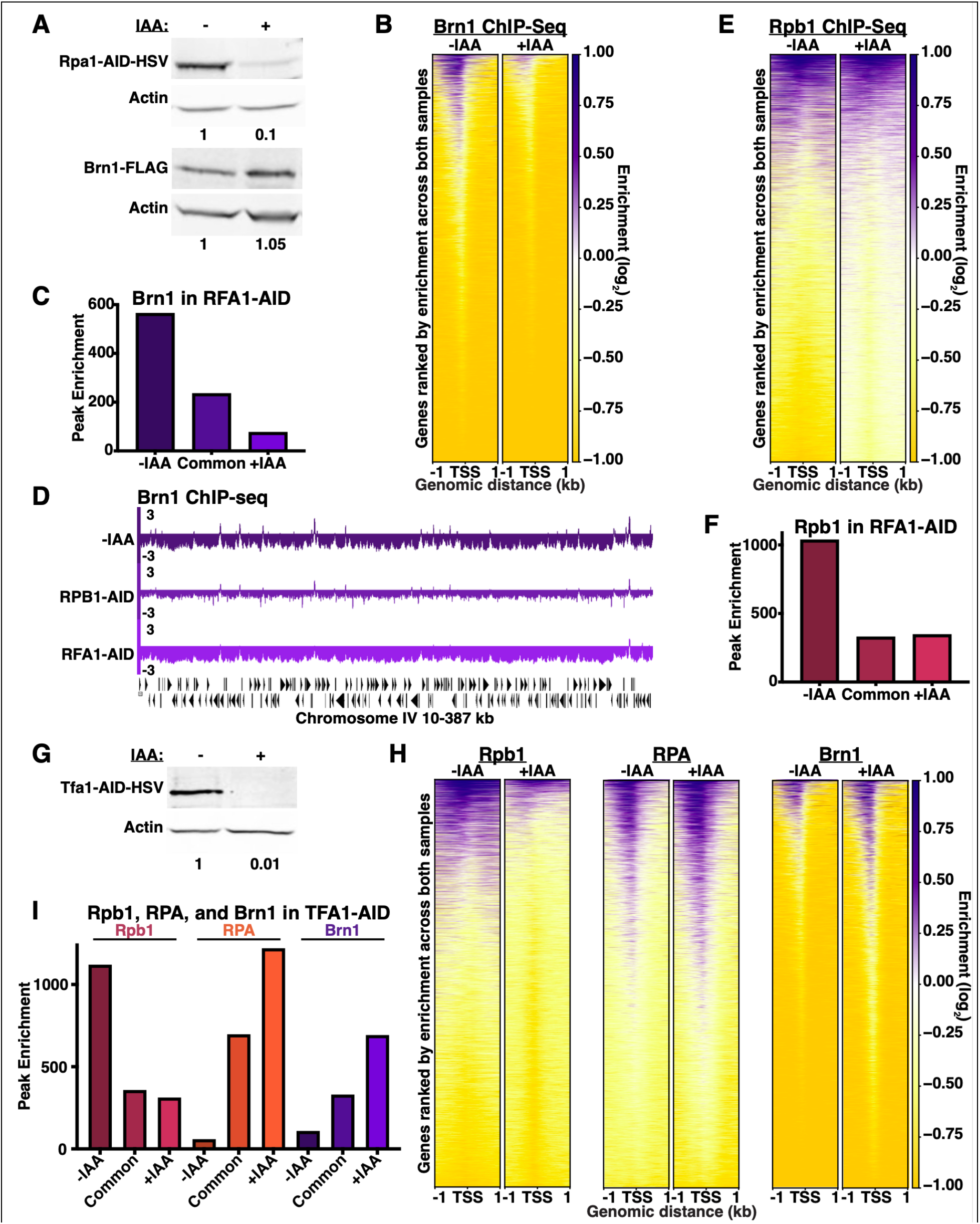
RPA, not transcription, is required for condensin targeting to promoters. (**A**) Representative Western blot in *RFA1*-AID-HSV Day 7 cells +/- IAA. Bands were normalized to actin and then to -IAA. (**B**) ChIP-seq heatmaps of Brn1-FLAG in *RFA1*-AID-HSV Day 7 cells +/- IAA. (**C**) Differential peak analysis of Brn1-FLAG in *RFA1*-AID-HSV Day 7 cells +/- IAA. (**D**) Representative genome browser view of ChIP-seq data. (**E**) ChIP-seq heatmaps of Rpb1 in *RFA1*-AID-HSV Day 7 cells +/- IAA. (**F**) Differential peak analysis of Rpb1 in *RFA1*-AID-HSV Day 7 cells +/- IAA. (**G**) Representative Western blot in TFA1-AID-HSV Day 7 cells +/- IAA. Bands were normalized to actin and then to -IAA. (**H**) ChIP-seq heatmaps of Rpb1, RPA, and Brn1 in TFA1-AID-HSV Day 7 cells +/- IAA. (**I**) Differential peak analysis of Rpb1, Brn1-FLAG, and RPA in TFA1-AID-HSV Day 7 cells +/- IAA.

Rpa1 depletion leads to a strong reduction in Brn1 signal (**Fig. 5B**). Differential peak calling and examination at the genome browser level reveals that most Brn1 peaks are lost when Rpa1 is depleted, and that this loss is even more pronounced than in Rpb1-depleted cells (**Fig. 5C-D**). To confirm these results, we depleted a different subunit of RPA, Rpa2, encoded by the *RFA2* gene (**Fig. S3A**). Consistently, Rpa2 depletion also results in a decrease in Brn1 ChIP-seq signal (**Fig. S3B-C**), though the effect is slightly weaker than for Rpa1 depletion (**Fig. S3D**), which we attribute to Rpa1 being the primary RPA subunit. We conclude that RPA is required for condensin localization at promoters during quiescence.

To determine that topological stress resulting from transcription and not transcription itself targets condensin, we performed ChIP-seq of Rpb1 in the *RFA1*-AID strains at Day 7. In the absence of Rpa1, Rpb1 occupancy also decreases (**Fig. 5E-F**), leaving us unable to attribute the decrease in condensin occupancy upon Rpa1 depletion solely to loss of RPA. To separate the contributions of RPA and transcription, we leveraged a system that has previously been used to shut down transcription, depletion of Tfa1, the largest subunit of TFIIE. As expected, IAA treatment during quiescence entry leads to loss of Tfa1 protein and Rpb1 occupancy in *TFA1*-AID strains at Day 7 (**Fig. 5G-I**). However, ChIP-seq of RPA and Brn1 shows that in the absence of Tfa1, both RPA and Brn1 levels increase at promoters, despite the loss of Rpb1 (**Fig. 5H-I**). We speculate that loss of TFIIE during quiescence may generate increased torsional stress at promoters and/or may lead to the formation of unstable transcriptional complexes that trigger the DNA damage response and ssDNA expansion.^82,83^ Regardless of the mechanism, Tfa1 depletion allows us to separate the roles of RNAPII and RPA on condensin localization. We conclude that transcription targets condensin indirectly during quiescence by causing RPA accumulation at promoters, and that it is RPA that directs condensin localization during quiescence.

### RPA does not target condensin in cycling cells

We next examined if RPA also targets condensin in cycling cells. We found that during cycling, RPA strongly overlaps with Brn1 at centromeres and tRNA genes, but not at the rDNA (**Fig. S4A-D**). We next depleted Rpa1 in cycling cells (**Fig. S4E**). Loss of RPA leads to a modest increase in Brn1 occupancy genome-wide, including at centromeres and tRNA genes (**Fig. S4F-H**). These results suggest that during the cell cycle, RPA not only does not target, but in fact antagonizes, condensin. This observation is consistent with reports that condensin overlaps with R-loops at tRNA genes in cycling cells, but resolution of R-loops does not affect condensin occupancy.^84^ Instead, location-specific factors such as TFIIIC, Sgo1, and Lrs4 likely target condensin in cycling cells.^85,86^

### RPA does not promote condensin loading in quiescent cells

In quiescent cells, RPA could direct condensin to active gene promoters by targeting condensin loading at these sites, by serving as a loop extrusion barrier, or both. To determine if RPA plays a role in condensin loading, we performed chromatin enrichment of Brn1-FLAG in Day 7 *RFA1*-AID cells (**Fig. 6A**). Surprisingly, we found an approximately 2-fold increase in the amount of chromatin-bound Brn1 in the absence of Rpa1. This suggests that rather than facilitating condensin loading, RPA limits condensin occupancy, perhaps by inhibiting loading or by contributing to condensin turnover. Consequently, RPA must target condensin by serving as a loop extrusion barrier during quiescence; when Rpa1 is depleted, ChIP-seq peaks decrease as condensin no longer accumulates at promoters and instead spreads throughout the genome. These results are consistent with the increase in condensin observed upon Rpa1 depletion in cycling cells (**Fig. S4F-H**), supporting that RPA antagonizes condensin occupancy in diverse locations and cell states.

**Figure 6.**
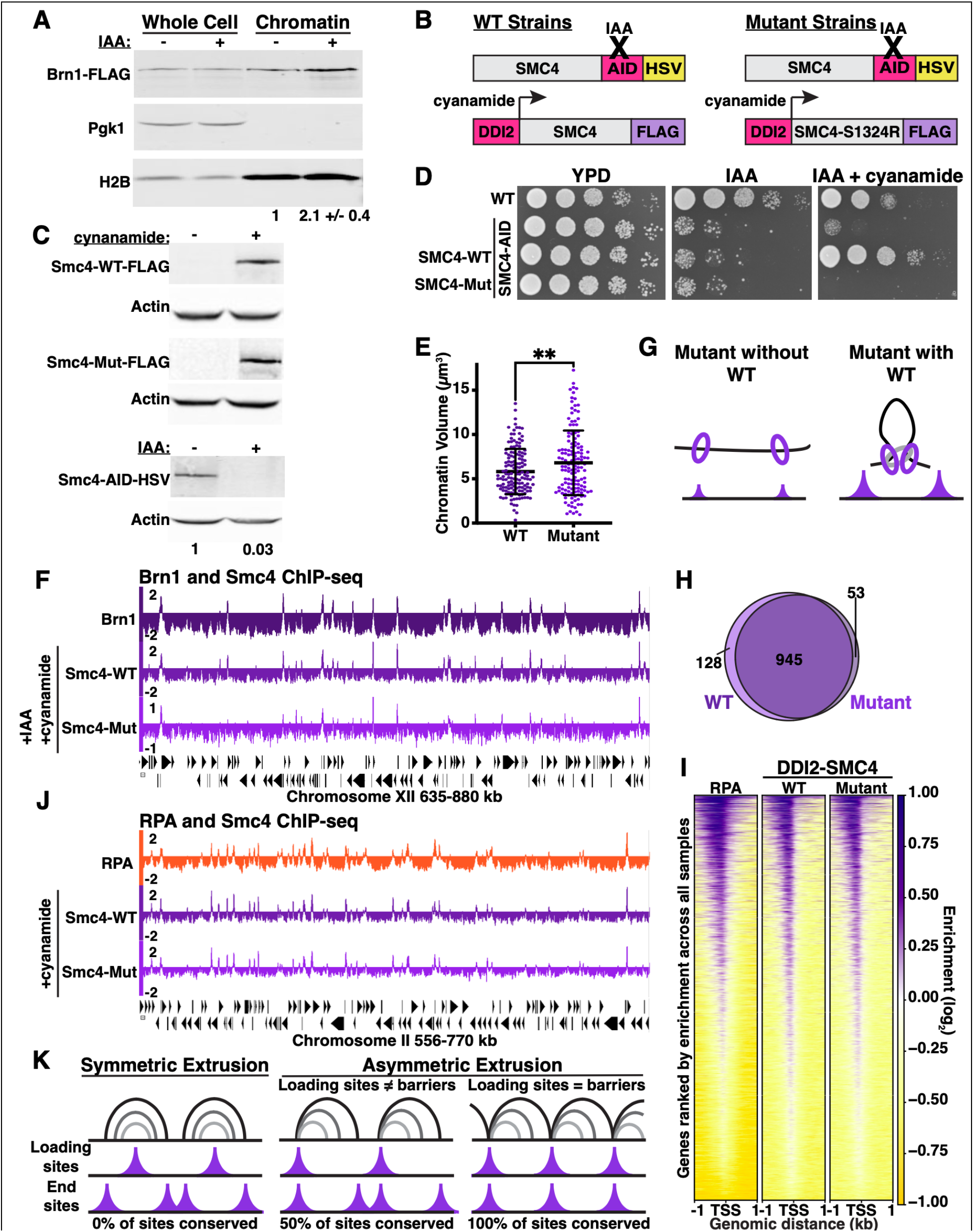
RPA sites are loading sites and extrusion barriers for asymmetric condensin loop extrusion. (**A**) Representative Western blot of yeast chromatin enriched fractions in *RFA1*-AID-HSV Day 7 cells +/- IAA. Pgk1 is a cytoplasmic control. Bands were normalized to H2B and then to -IAA. Numbers are the mean and SD of biological duplicates. (**B**) Schematic of *SMC4*-WT and *SMC4*-Mut strains. (**C**) Representative Western blot of Day 7 *SMC4*-WT and *SMC4*-Mut strains +/- cyanamide and IAA. For Smc4-AID-HSV, band was normalized to actin and then to -IAA. (**D**) Spot assay of WT, SMC4-AID, SMC4-WT, and SMC4-Mut strains +/- cyanamide and IAA. (**E**) Chromatin volume quantification of Z-stacks of SMC4-WT and SMC4-Mut Day 7 cells treated with IAA and cyanamide. Bars represent mean +/- SD. Significance measured using unpaired Welch’s t-test. (**F**) Representative genome browser view of ChIP-seq data. (**G**) Schematic of Smc4-mut in the absence and presence of untagged WT endogenous Smc4. (**H**) Direct overlap of ChIP-seq peaks. Peaks were filtered using a 1.5-fold enrichment threshold. (**I**) ChIPseq heatmaps of RPA, Smc4-WT, and Smc4-Mut in Day 7 cells + cyanamide in the presence of endogenous Smc4. (**J**) Representative genome browser view of ChIP-seq data. (K) Schematic of loop extrusion progression and the resulting ChIP-seq peaks for symmetric and asymmetric extrusion.

### RPA-coated ssDNA sites are both loading sites and extrusion barriers for condensin

We next sought to identify condensin loading sites in our system. To this end, we took advantage of a previously reported *SMC4* ATPase mutation, *SMC4-S1324R*.^87^ Smc4 bearing this mutation is incorporated into condensin complexes, but while the resulting complexes remain able to load onto chromatin, they cannot hydrolyze ATP or extrude loops.^87^ Consequently, Smc4-S1324R binding sites are loading sites, whereas wild-type (**WT**) condensin binding sites are extrusion end sites that could be loading sites, extrusion barriers, or both. We constructed strains in which the endogenous *SMC4* allele is tagged with AID and HSV, and WT (***SMC4*-WT**) or S1324R (***SMC4*-Mut**) copies tagged with FLAG under control of the *DDI2* promoter are integrated at the *URA3* locus (**Fig. 6B**). Addition of cyanamide to these strains induces expression of the exogenous *SMC4* copies to similar levels, while addition of IAA depletes the endogenous protein (**Fig. 6C**). To test the functionality of these strains, we completed a yeast spot assay. In the presence of IAA, *SMC4*-WT and *SMC4*-Mut strains display similar growth defects, consistent with the essentiality of *SMC4* (**Fig. 6D**). However, in the presence of IAA and cyanamide, only *SMC4*-WT rescues depletion of endogenous Smc4. To test the ability of these strains to condense chromatin, we additionally tagged H2B with GFP and measured chromatin volume in IAA and cyanamide-treated Day 7 cells via confocal microscopy (**Fig. 6E**). *SMC4*-Mut cells display a decrease in chromatin condensation compared to *SMC4*-WT, consistent with *SMC4-S1324R*’s previously reported condensation defect in mitotic cells.^87^ To map condensin loading sites, we completed ChIP-seq against FLAG in Day 7 *SMC4*-WT and *SMC4*-Mut cells treated with IAA and cyanamide during quiescence entry and compared the data to Brn1-FLAG in a WT background (**Fig. 6F**). The locations of ChIP-seq peaks across these samples are virtually identical. This suggests that most condensin binding sites are both loading sites and extrusion barriers.

Our next goal was to quantify the overlap between *SMC4*-WT and *SMC4*-Mut and to determine their colocalization with RPA. Although the pattern of Smc4-Mut binding is nearly identical to Smc4-WT, the ChIP signals are markedly lower in *SMC4*-Mut (**Fig. 6F**), making peak calling challenging. We reasoned that the topology of barrier sites that have blocked condensin may facilitate condensin crosslinking and antibody capture, leading to enhanced ChIP signal. Therefore, we repeated ChIP-seq in *SMC4*-WT and *SMC4*-Mut strains in a WT background in the presence of cyanamide (**Fig. 6G**). The presence of untagged endogenous Smc4 leads to Smc4-Mut signals equivalent to Smc4-WT (**Fig. 6H-J**), confirming our hypothesis that loading sites are detected by ChIP less efficiently when they do not additionally function as extrusion barriers. Additionally, Smc4-Mut and Smc4-WT overlap at the vast majority of locations, consistent with loading sites also serving as extrusion barriers (**Fig. 6 H-J**). Comparison of these results to RPA Day 7 ChIP-seq data shows near complete overlap between condensin loading sites, end sites, and RPA (**Fig. 6I-J**). We conclude that although RPA does not promote condensin loading, RPA sites serve both as condensin loading sites and extrusion barriers in quiescent cells.

### Condensin extrudes loops asymmetrically in quiescent yeast cells

SMC complexes have been proposed to extrude loops either symmetrically or asymmetrically.^23,25,26^ During symmetric extrusion, complexes load onto chromatin and extrude loops in both directions until reaching extrusion barriers on either side, resulting in end sites that are distinct from loading sites (**Fig. 6K**).^25,26^ During asymmetric extrusion, complexes load onto chromatin and extrude loops in only one direction, resulting in 50% conservation of loading sites at end sites.^23,26^ Complete conservation of loading and end sites, demonstrated by the overlap between Smc4-WT and Smc4-Mut, suggests asymmetric extrusion in which loading sites and extrusion barriers are the same. Consequently, we deduce that in quiescent yeast cells, condensin extrudes loops asymmetrically, consistent with single-molecule studies of purified yeast condensin.^23^

### Loss of RPA leads to large-scale chromatin reorganization

We next sought to establish the role of RPA in regulating condensin-mediated 3D chromatin architecture. To do so, we constructed H2B-GFP strains in *RFA1*-AID and *SMC4*-AID backgrounds and measured chromatin volume by confocal microscopy in Day 7 cells (**Fig. S5A**). Loss of Smc4 leads to a decrease in chromatin condensation similar to *SMC4*-Mut cells (**Fig. S5B** and **Fig. 6D**). Despite more condensin being present on the genome than in -IAA cells, Rpa1 depletion leads to a similar reduction in chromatin condensation as Smc4 depletion (**Fig. S5A-B**), suggesting condensin function may be altered in these cells.

To examine changes in 3D structure in *RFA1*-AID cells in detail, we performed Micro-C in Day 7 cells in biological duplicate in the presence and absence of IAA. After confirming agreement between replicates,^88^ we merged replicate data to maximize signal to noise (**Fig. S5C**). In budding yeast, centromere clustering at the spindle pole body produces X-shaped contact patterns of interchromosomal interactions between regions of chromatin within 200 kilobases (**kb**) of centromeres.^60,89^ This clustering increases in quiescent cells in which Smc4 is depleted.^60^ In contrast, Rpa1-depleted cells display loss centromere clustering in genome-wide contact maps and pileup plots (**Fig. S5D-F**), consistent with an increase in condensin levels. These results suggest that loss of RPA leads to changes in condensin-mediated chromatin condensation at the global level.

### RPA regulates condensin loop extrusion dynamics

To decipher the role of RPA in condensin loop extrusion, we next analyzed the Micro-C data at high resolution. Contact maps at 1 kb and 200 base-pair (**bp**) resolution reveal a striking increase in striping in the absence of RPA (**Fig. 7A-C**). Stripes are believed to arise from loop extrusion, during which regions of chromatin are transiently brought together by the SMC complex during loop expansion.^26,90^ Closer examination reveals that stripes emanate from the same locations + and -IAA (**Fig 7A**), suggesting condensin loading sites are preserved in the absence of RPA. Importantly, the pattern of striping observed in both samples is consistent with asymmetric extrusion. Stripes associated with asymmetric extrusion extend from the diagonal at a 45° angle, in contrast to perpendicular stripes associated with symmetric extrusion.^26,91,92^ In this case, stripes begin and end at the same sites, consistent with loading sites also being extrusion barriers. The stripes also form closed right triangles, further suggesting that condensin molecules may load at boundaries on each end and extrude loops in opposite directions, either at the population level or within the same cells. Although stripes are present in both the -IAA and +IAA samples, stripes are more prominent and span substantially greater distance upon loss of RPA (**Fig. 7A-C**). The increase in the distance of stripes in the +IAA sample suggests that though loading sites are preserved, barrier function is disrupted when RPA is depleted.

**Figure 7:**
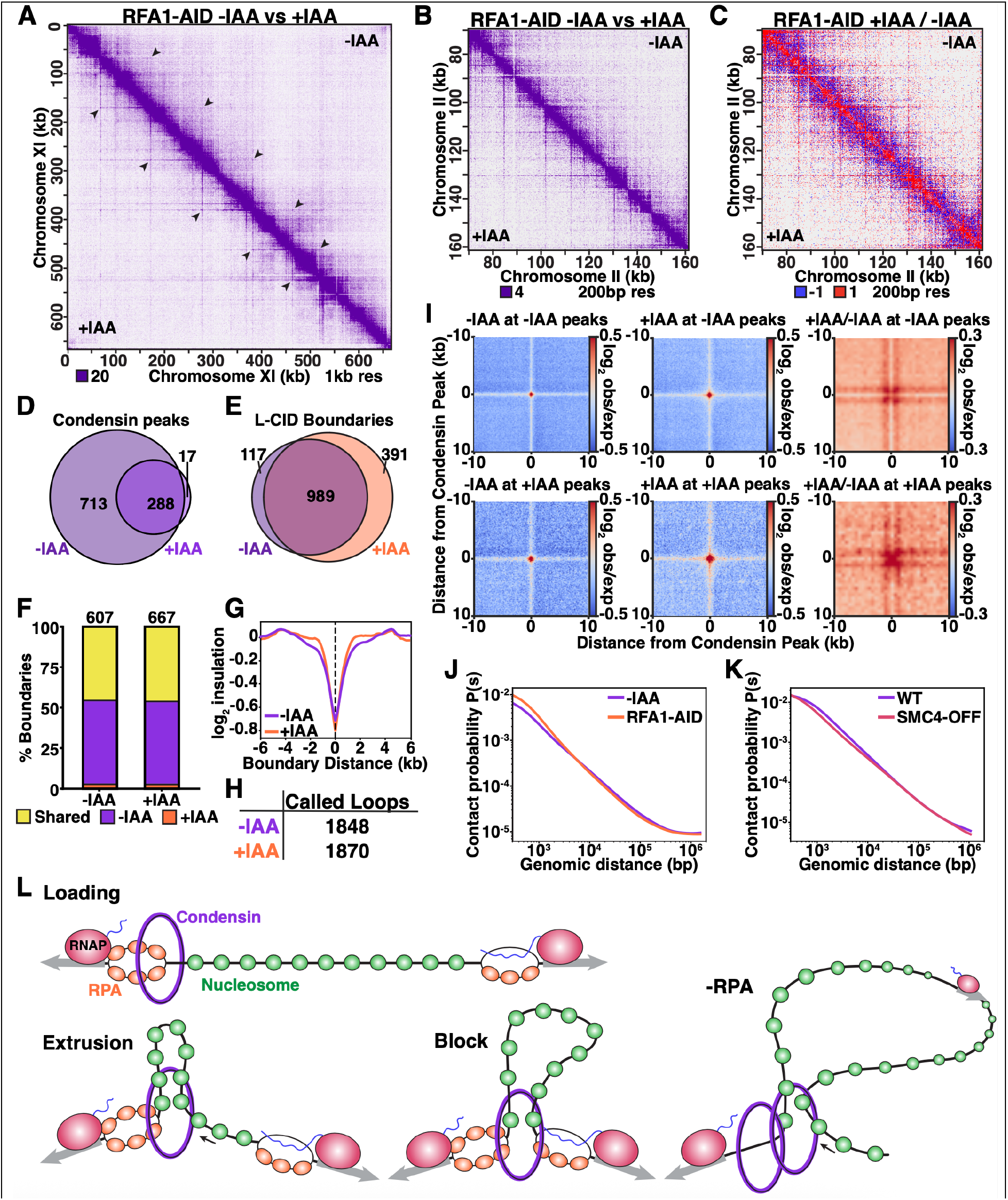
RPA regulates condensin loop extrusion. (**A**) Micro-C contact map in *RFA1*-AID Day 7 cells +/- IAA. Arrows point to stripes in the same locations +/-IAA treatment. (**B**) Micro-C contact map in *RFA1*-AID Day 7 cells +/- IAA. (**C**) Micro-C contact map in *RFA1*-AID Day 7 cells. +IAA data are divided by -IAA data. (**D**) Direct overlap of Brn1-FLAG ChIP-seq peaks filtered using a 1.5-fold enrichment threshold. (**E**) Overlap of called L-CID boundaries in *RFA1*-AID Day 7 cells +/- IAA. (**F**) Percent L-CID boundaries overlapping with Brn1 peaks. (**G**) Insulation plots centered at L-CID boundaries overlapping shared Brn1 peaks. (**H**) Number of called loops in *RFA1*-AID Day 7 cells +/- IAA. (**I**) Pileup plots of 150 bp Micro-C data centered on Brn1 peaks in the indicated condition. Right plots show +IAA data divided by -IAA data. (**J**) Contact probability of *RFA1*-AID Day 7 cells +/- IAA. (**K**) Contact probability of WT and SMC4-off Day 7 cells. Data from Swygert, et al., 2019. (**L**) Model. See text for details.

Although condensin loading sites did not appear altered in the Micro-C contact maps, we quantified Brn1 ChIP-seq peaks and calculated their overlap independent of peak enrichment to determine if new condensin sites arise upon RPA depletion (**Fig. 7D**). Nearly all Brn1 peaks in the +IAA sample directly overlap with peaks in the -IAA sample, suggesting condensin does not load at new sites in the absence of RPA, though we cannot rule out dispersed loading at additional sites that dilute out at the population level. We next called L-CID boundaries in each sample and determined their overlap with condensin peaks present in either -IAA, +IAA, or both (**Fig. 7E-F**). Although slightly more boundaries could be called in the +IAA sample (**Fig. 7E**), most new boundaries do not overlap with condensin peaks in either -IAA or +IAA, and, overall, boundaries overlapping with condensin +/-IAA are conserved across samples (**Fig. 7F**). Despite the decrease in condensin ChIP-seq peaks, boundaries overlapping Brn1 peaks present in -IAA, +IAA, or both are all more insulating in the absence of RPA (**Fig. 7G** and **Fig. S6A**). The number of called loops is also very similar across samples (**Fig. 7H**). These results are in stark contrast to our previous results in Day 7 cells in which Smc4 is depleted via a tetracycline-off system (***SMC4*-off**), in which half of all boundaries are lost, the insulation of remaining boundaries decreases, and looping is greatly reduced.^60^ Altogether, these results suggest that condensin loading sites are maintained in the absence of RPA, even at locations where Brn1 cannot be detected by ChIP-seq in the +IAA sample.

As postulated above, if condensin loads at the same sites in the presence and absence of RPA, the decrease in Brn1 ChIP-seq signal in the +IAA sample necessarily points to the loss of extrusion barrier function at RPA sites, consistent with the increased distance of striping observed in the contact maps (**Fig. 7A-C**). To confirm this genome-wide, we generated pileup plots centered on condensin peaks present in either - or +IAA (**Fig. 7I**). These plots reveal stronger striping in the +IAA sample at both sets of condensin peaks. In addition to being stronger, stripes are also wider in the absence of RPA, consistent with the loss of extrusion barriers resulting in less positioned loops. This finding is supported by pileup plots at loops in which at least one loop anchor overlaps with a condensin site (**Fig. S6B**).

To determine how these altered extrusion dynamics affect chromatin folding, we calculated how contact probability is changing relative to genomic distance (**Fig. 7J**). In +IAA samples, local interactions up to ∼6 kb increase, consistent with increased condensin occupancy. Beyond 6 kb, interactions decrease, suggesting L-CID loops, which range from 1-60 kb and average around 13 kb,^60^ are less stable absent RPA extrusion barriers. These results are in direct contrast to changes in contact probability in *SMC4*-off cells, in which local contacts decrease up to around 20 kb (**Fig. 7K**). To confirm this result, we generated on-diagonal pileup plots centered at condensin peaks (**Fig. S6C**). Although interactions within loops increase in the +IAA samples up to ∼10 kb, interactions within loops beyond this point are higher in the -IAA sample. These pattens of contacts can also be seen in the differential contact map, in which very local contacts and stripes are higher in +IAA, but the majority of contacts between stripes (i.e., within loops) are higher in -IAA (**Fig. 7C**). These results are consistent with a model in which more condensin is present on chromatin in the absence of RPA, but disruption of extrusion barrier function leads to fewer contacts within loops.

## DISCUSSION

Based on these findings and our previous observation that condensin localization in quiescent cells correlates with nucleosome depleted regions (**NDR**s),^64,93^ we propose the following model (**Fig. 7L**). During quiescence entry, transcription in the absence of topoisomerases generates RPA-coated ssDNA at NDRs in promoters. Condensin loads at these sites and extrudes loops asymmetrically until reaching a subsequent RPA site, gradually accumulating at promoter regions over the course of several days. Once stabilized at RPA extrusion barriers, condensin loops promote interactions between nucleosomes within the loop, further repressing transcription in quiescent cells.^60^ In the absence of RPA, condensin occupancy at promoter NDR regions increases, increasing local contacts between nucleosomes near loading sites. Condensin then continues elongating loops indefinitely, preventing the loop stabilization that promotes longer-range interactions within loops. Transcription also decreases upon loss of RPA, perhaps because RPA-bound structures contribute to maintaining open promoters. Alternately, in the absence of extrusion barriers at transcribed gene promoters, condensin molecules extrude through genes and disrupt transcription, similar to what has been recently proposed for cohesin loaded near genes.^44^

A plethora of evidence supports an antagonistic relationship between condensin and RPA. Condensin has repeatedly been found to induce positive supercoiling and to bind ssDNA, double-stranded DNA (**dsDNA**), and supercoiled DNA *in vitro*.^23,80,94–99^ Further, condensin is well-documented to cooperate with topoisomerases to relieve torsional stress during mitotic chromosome condensation and replication stress.^100–103^ A key biochemical study demonstrated that condensin can bind RPA-coated ssDNA and induce positive supercoiling that reanneals DNA and evicts RPA, which was later suggested also to occur in mitotic fission yeast.^80,81^ Further, RPA mutants suppress mutations in the condensin hinge region, which selectively binds ssDNA.^80,97,99,104^ Condensin has also been shown to co-localize with R-loops in cycling cells.^84^

Beyond condensin, RPA and ssDNA have been implicated in regulating the function of cohesin and SMC5/6 in biochemical assays.^19,105–109^ SMC5/6 favors binding to ssDNA/dsDNA junctions, which is inhibited by RPA.^107–109^ Similarly, cohesin can simultaneously bind ssDNA and dsDNA, with ssDNA binding inhibited by RPA.^105^ Loop extrusion by human cohesin is stalled by R-loops in single-molecule assays, and cohesin STAG1 and STAG2 subunits bind R-loops *in vitro*.^110,111^ Correlative evidence also links cohesin to ssDNA in mammalian cells, as cohesin ChIP-seq peaks overlap with both R-loops and G4s, particularly in the absence of CTCF.^111,112^

Here, we demonstrate that most condensin sites overlap with RPA both in cycling and quiescent budding yeast cells, and that loss of RPA leads to an increase in condensin levels on the genome in both cell states. RPA-bound sites of transcription-induced topological stress, including R-loops and G4s, further serve to block condensin loop extrusion during quiescence. These results connect disparate results from biochemical studies into a model based on direct evidence in cells, uncover the relationship between transcription and condensin targeting, and suggest antagonism by RPA may be a universal driver of SMC complex function.

We also show that condensin extrudes loops asymmetrically in cells, consistent with predictions made by biochemical single-molecule assays in which yeast condensin exclusively performs asymmetrical, one-sided extrusion.^23,113^ Interestingly, modeling studies suggest that one-sided asymmetric extrusion can only produce striping patterns in contact maps and efficiently compact the genome if extruders load at defined locations.^26^ These results are entirely concordant with our findings that condensin loads specifically at RPA sites. That loading sites are also extrusion barriers, with extrusion occurring bidirectionally from each site, provides an efficient mechanism to eliminate gaps in chromatin compaction, generating bottle brush structures of loops separated by boundaries at transcribed genes.

How does RPA-coated ssDNA antagonize SMC complex occupancy and extrusion? The simplest possibility is that SMC complexes compete with RPA for ssDNA binding. Alternately, RPA nucleoprotein filaments may be too stiff to be extruded.^114,115^ RPA filaments have also been shown to phase separate,^116,117^ perhaps generating condensates that are impermeable by SMC complexes. Ultimately, understanding the underlying mechanism will require further investigation.

### Limitations of the study

The primary limitation of this study is that we do not know what happens to the underlying DNA when we deplete RPA. RPA depletion in proliferating cells results in the accumulation of DNA double-strand breaks and replication catastrophe. However, the need for RPA is likely reduced in quiescent cells that do no undergo replication. We do not see evidence of widespread DNA damage in sequenced ChIP input DNA, microscopy, or Micro-C data in the absence of RPA, and we are able to obtain a robust population of Rpa1-depleted quiescent cells for use in experiments. We speculate that upon RPA depletion in quiescent cells, some DNA structures may reanneal while others persist as ssDNA. A secondary limitation is that we present several lines of evidence that ChIP-seq may be less efficient at capturing condensin loading sites than barrier sites when the two are separated. An alternate method may be more suitable for specifically capturing loading.

## RESOURCE AVAILABILITY

## Lead contact

Requests for further information and resources should be directed to and will be fulfilled by the lead contact, Sarah G. Swygert.

## Materials availability

All yeast strains and plasmids generated by this study are available from the lead contact without restriction.

## Data and code availability

### Data

All ChIP-seq and Micro-C XL data will be deposited at NCBI GEO and made publicly available as of the date of publication.

### Code

The Micro-C XL analysis pipeline can be found on the SwygertLab GitHub page at: https://github.com/SwygertLab/Micro-C

### Additional information

Any additional information required to reanalyze the data reported in this paper is available from the lead contact upon request.

## ACKNOWLEDGMENTS

The authors would like to thank Toshio Tsukiyama, Laurie Stargell, and members of the Swygert lab for insightful discussion. Research was supported by National Institutes of Health grants R00GM134150 and R35GM154898 to S.G.S., R35GM139483 to S.M.M., and a Schlumberger Faculty of the Future Fellowship to B.A. This study was supported in part by the NIH P30CA046934 funded Genomics Shared Resource Core Facility (RRID SCR_021984). This work also utilized the Alpine High-Performance Computing resource, which is funded by the University of Colorado Boulder, the University of Colorado Anschutz, Colorado State University, and the National Science Foundation (award 2201538). Fig. 3D was created using Biorender.

## AUTHOR CONTRIBUTIONS

B.A. and S.G.S. designed the study, performed and analyzed genomics experiments, and wrote the first draft of the manuscript. J.A.H. and B.A developed analysis pipelines. B.A., A.H.L., L.M.S., and S.G.S. constructed strains and performed genetics and molecular biology assays. S.M.M. supervised the microscopy experiments, A.H.L. optimized slide preparation, B.A. and S.M.M. performed microscopy, and B.A. analyzed the microscopy data. S.G.S. supervised B.A., J.A.H., A.H.L., and L.M.S. All authors assisted in editing the manuscript.

## SUPPLEMENTAL INFORMATION

Figures S1-S6.

## STAR METHODS

## KEY RESOURCES TABLE

| REAGENT or RESOURCE | SOURCE | IDENTIFIER |
| --- | --- | --- |
| **Antibodies** |  |  |
| Anti-Rpb1 (mouse monoclonal)<br>2 µl for ChIP | Cell signaling | #2629 |
| Anti-FLAG M2 (mouse monoclonal)<br>4 µl for ChIP, 1:2,500 to 1:5,000 for WB | Sigma | #F1804 |
| Anit-RPA (rabbit polyclonal)<br>4 µl for ChIP, 1:5,000 for WB | Agrisera | #AS07 214 |
| Anti-H3 (rabbit polyclonal)<br>1 µl for ChIP | Abcam | #1791 |
| Anti-HSV (rabbit polyclonal)<br>1:5,000 for WB | Sigma | #H6030 |
| Anit-H2B (rabbit polyclonal)<br>1:5,000 for WB | Active Motif | #39237 |
| Anti-Actin (mouse monoclonal)<br>1:20,000 for WB | Abcam | #8224 |
| Anti-PGK1 (rabbit polyclonal)<br>1:500 for WB | Invitrogen | #PA5-28612 |
| **Chemicals, peptides, and recombinant proteins** |  |  |
| Disuccinimidyl glutarate (DSG) | Thermo Fisher Scientific | #PI20593 |
| Indole-3-acetic acid (IAA) | Sigma | #I3750-5G-A |
| Percoll | Cytiva | #17-0891-01 |
| Cyanamide | Sigma | #C87908 |
| Micrococcal nuclease | Worthington | #LS004798 |
| Zymolyase 100T | amsbio | #120493-1 |
| **Critical commercial assays** |  |  |
| NEBNext® Ultra™ II DNA Library Prep Kit for Illumina | NEB | #E7645 |
| Ovation Ultralow V2 Library kit | NuGen | #0344 |
| **Deposited data** |  |  |
| Micro-C | This paper | GEO accession # to be added |
| ChIP-Seq | This paper | GEO accession # to be added |
| **Experimental models: Organisms/strains** |  |  |
| *S. cerevisiae*: W303 MATa RAD5+ | McKnight et al., 2015 | yTT5781 |
| *S. cerevisiae*: W303 MATa RAD5+ | McKnight et al., 2015 | yTT5783 |
| *S. cerevisiae*: W303 MATa RAD5+ BRN1-2L-3xFLAG-KanMX | This paper | ySGS80 |
| *S. cerevisiae*: W303 MATa RAD5+ BRN1-2L-3xFLAG-KanMX | This paper | ySGS81 |
| *S. cerevisiae*: MATa RAD5+ HTB2-GFP-KanMX | This paper | ySGS186 |
| *S. cerevisiae*: MATa RAD5+ HTB2-GFP-KanMX | This paper | ySGS187 |
| *S. cerevisiae*: W303 MATa RAD5+ TOP1-2L-3xFLAG-KanMX | This paper | ySGS177 |
| *S. cerevisiae*: W303 MATa RAD5+ TOP1-2L-3xFLAG-KanMX | This paper | ySGS178 |
| *S. cerevisiae*: W303 MATa RAD5+ TOP2-2L-3xFLAG-KanMX | This paper | ySGS179 |
| *S. cerevisiae*: W303 MATa RAD5+ TOP2-2L-3xFLAG-KanMX | This paper | ySGS180 |
| *S. cerevisiae*: W303 MATa can1-100 RAD5+ BRN1-FLAG-KanMX URA3-1::ADH1-OsTIR RPB1-HSV-AID-Hyg | This paper | ySGS511 |
| *S. cerevisiae*: W303 MATa can1-100 RAD5+ BRN1-FLAG-KanMX URA3-1::ADH1-OsTIR RPB1-HSV-AID-Hyg | This paper | ySGS513 |
| *S. cerevisiae*: W303 MATa can1-100 RAD5+ BRN1-FLAG-KanMX URA3-1::Adh1-OsTIR TFA1-HSV-AID-Hyg | This paper | ySGS515 |
| S. cerevisiae: W303 MATa can1-100 RAD5+ BRN1-FLAG-KanMX URA3-1::Adh1-OsTIR TFA1-HSV-AID-Hyg | This paper | ySGS517 |
| S. cerevisiae: W303 MATa can1-100 RAD5+ BRN1-FLAG-KanMX URA3-1::ADH1-OsTIR RFA1-HSV-AID-Hyg | This paper | ySGS562 |
| S. cerevisiae: W303 MATa can1-100 RAD5+ BRN1-FLAG-KanMX URA3-1::ADH1-OsTIR RFA1-HSV-AID-Hyg | This paper | ySGS564 |
| S. cerevisiae: W303 MATa can1-100 RAD5+ Brn1-FLAG-KanMX URA3-1::ADH1-OsTIR RFA2-HSV-AID-Hyg | This paper | ySGS566 |
| S. cerevisiae: W303 MATa can1-100 RAD5+ Brn1-FLAG-KanMX URA3-1::ADH1-OsTIR RFA2-HSV-AID-Hyg | This paper | ySGS568 |
| S. cerevisiae: W303 MATa can1-100 RAD5+ BRN1-FLAG-KanMX URA3-1::ADH1-OsTIR SMC4-HSV-AID-Hyg | This paper | ySGS549 |
| S. cerevisiae: W303 MATa can1-100 RAD5+ BRN1-FLAG-KanMX URA3-1::ADH1-OsTIR SMC4-HSV-AID-Hyg | This paper | ySGS551 |
| S. cerevisiae: W303 MATa W303 RAD5+ can1-100 his3-11 SMC4-HSV-AID-Hyg SSN6:: ADH1-OsTIR-HIS URA3::DDI2-SMC4WT | This paper | ySGS555 |
| S. cerevisiae: W303 MATa W303 RAD5+ can1-100 his3-11 SMC4-HSV-AID-Hyg SSN6:: ADH1-OsTIR-HIS URA3::DDI2-SMC4WT | This paper | ySGS557 |
| S. cerevisiae: W303 Mata W303 RAD5+ can1-100 his3-11 SMC4-HSV-AID-Hyg SSN6:: ADH1-OsTIR-HIS URA3::DDI2-SMC4S1324R | This paper | ySGS559 |
| S. cerevisiae: W303 Mata W303 RAD5+ can1-100 his3-11 SMC4-HSV-AID-Hyg SSN6:: ADH1-OsTIR-HIS URA3::DDI2-SMC4S1324R | This paper | ySGS561 |
| S. cerevisiae: W303 Mata W303 RAD5+ ura3-1 can1-100 his3-11 SMC4-HSV-AID-Hyg SSN6:: ADH1-OsTIR-HiS HTB2-GFP-KanMX | This paper | ySGS570 |
| S. cerevisiae: W303 Mata W303 RAD5+ ura3-1 can1-100 his3-11 SMC4-HSV-AID-Hyg SSN6:: ADH1-OsTIR-HiS HTB2-GFP-KanMX | This paper | ySGS572 |
| S. cerevisiae: W303 MATa Rad5+ can1-100 RFA1-HSV-AID-Hyg HTB2-GFP-KanMX URA3::ADH1-OsTIR | This paper | ySGS578 |
| S. cerevisiae: W303 MATa Rad5+ can1-100 RFA1-HSV-AID-Hyg HTB2-GFP-KanMX URA3::ADH1-OsTIR | This paper | ySGS580 |
| S. cerevisiae: W303 Mata RAD5+ can1-100 his3-11 SMC4-HSV-AID-Hyg SSN6::ADH1-OsTIR-HiS URA3::DDI2-SMC4WT HTB2-GFP-KanMX | This paper | ySGS588 |
| S. cerevisiae: W303 Mat@ RAD5+ can1-100 his3-11 SMC4-HSV-AID-Hyg SSN6::ADH1-OsTIR-HiS URA3::DDI2-SMC4WT HTB2-GFP-KanMX | This paper | ySGS590 |
| S. cerevisiae: W303 Mata W303 prototroph RAD5+ can1-100 his3-11 SMC4-HSV-AID-Hyg SSN6::ADH1-OsTIR-HiS URA3::DDI2-SMC4S1324R HTB2-GFP-KanMX | This paper | ySGS592 |
| S. cerevisiae: W303 Mat@ W303 prototroph RAD5+ can1-100 his3-11 SMC4-HSV-AID-Hyg SSN6::ADH1-OsTIR-HiS URA3::DDI2-SMC4S1324R HTB2-GFP-KanMX | This paper | ySGS593 |
| S. cerevisiae: W303 MATa can1-100 RAD5+ URA3::DDI2-RNH1D193N-2xL-3xFLAG | This paper | ySGS504 |
| S. cerevisiae: W303 MATa can1-100 RAD5+ URA3::DDI2-RNH1D193N-2xL-3xFLAG | This paper | ySGS506 |
| **Software and algorithms** |  |  |
| bowtie2 | Langmead and Salzberg, 2012 | https://bowtie-bio.sourceforge.net/bowtie2/index.shtml |
| SAMtools | Li et al., 2009 | https://samtools.sourceforge.net/ |
| deepTools | Ramirez et al., 2016 | https://deeptools.readthedocs.io/en/latest/index.html |
| MACS2 | Zhang et al., 2008 | https://github.com/macs3-project/MACS |
| bedtools | Quinlan et al., 2010 | https://github.com/arq5x/bedtools2 |
| bwa-mem2 | Vasimuddin et al 2019 | https://github.com/bwa-mem2/bwa-mem2 |
| pairtools | Abdennur et al ., 2024 | https://github.com/open2c/pairtools |
| juicer | Durand et al., 2016 | https://github.com/aidenlab/juicer |
| cooltools | Abdennur et al., 2024 | https://github.com/open2c/cooltools |
| coolpuppy | Flyamer et al., 2020 | https://github.com/open2c/coolpuppy/blob/master/README.md |
| chromosight | Matthey-Doret et al., 2020 | https://github.com/koszullab/chromosight |
| hicrep | Yang et al., 2017 | https://github.com/dejunlin/hicrep |
| G4Hunter | Brazda et al., 2019 | https://github.com/AnimaTardeb/G4Hunter |

## METHOD DETAILS

### Yeast culture and quiescent cell purification

Prototrophic W303 *Saccharomyces cerevisiae* strains were constructed using standard techniques as detailed in the key resources table. Yeast were grown in YPD media (1% yeast extract, 2% Bacto Peptone, 2% glucose) with pH adjusted to 5.5 in a 30°C incubator with shaking at 200 rpm. Cell cultures were started from a single colony from a plate streaked no longer than five days prior. Quiescent cell purification was done as previously described.^51,118^ Briefly, on the day of purification 50 mL of culture was pelleted and resuspended in 1 mL of ultrapure water. Density gradients were prepared containing 90% (v/v) Percoll (Cytiva, catalog #17-0891-01) and 150 mM NaCl. Gradient mixtures were centrifuged at 10,000 rcf for 15 minutes at 4°C with slow acceleration and deceleration. Cell suspensions were layered over the gradient and centrifuged at 350 rcf for 2 hours at 4°C, and the bottom fraction of cells was isolated. Quiescent cells were washed with ultrapure water and cells were quantified using a spectrophotometer at 660 nm. For quiescence entry time courses, quiescent cells were purified as above on the indicated day, with the day the culture was started designated as Day 0. Cycling cells were grown to optical density (**OD**) 0.8-1 using standard methods.

### Transcriptional induction

Strains were constructed with FLAG-tagged genes under control of the inducible *DDI2* promoter as previously described.^79^ For *SMC4* induction, cyanamide (Sigma, catalog #C87908) was add to cultures to 18 mM final concentration on day 3 of quiescence entry. For RNH1-D193N expression, cyanamide was added to 18 mM on between day 6 and day 7 to achieve equivalent moderate levels of induction across replicates. Induction was verified by Western blot.

### Targeted protein degradation

Strains expressing OsTIR with target genes tagged with HSV and an auxin inducible degron were grown in 5 mL overnight cultures, which were used to start new 50 mL or 100 mL cultures to OD 0.0096.^119^ Cells were grown for ∼15 hours and glucose was monitored using glucose strips to determine the time of diauxic shift (glucose exhaustion). For all depletions except Rpb1, IAA (Sigma, catalog #I3750-5G-A) was added at 6 hours post diauxic shift as a powder at a final concentration of 1 mg/mL. To minimize the effect of Rpb1 depletion on condensin expression, IAA was added at 26 hours post diauxic shift. Quiescent cells were purified as detailed above and depletion was confirmed by Western blot. For cycling cells, cultures were started at OD 0.25 in 250 mL cultures and grown until reaching OD ∼1. IAA was added 30 minutes before cultured reached the desired density.

### Protein extract preparation

30 OD units of purified quiescent cells or 10 OD units of cycling cells were pelleted, washed with cold ultrapure water with 1x PMSF, flash frozen, and stored at −80°C. Trichloroacetic acid (**TCA**) protein extractions were performed as described with modification.^120^ Briefly each pellet was resuspended in 250 µL of 20% TCA and lysed by bead beating for 6-10 minutes. To collect cell lysates, a needle was used to make a hole in the bottom of the tube and tubes were placed in 15 mL conical tubes. Samples were centrifuged at 600 rpm for 2 minutes at 4°C. Beads were washed with 500 µl 5% TCA, and flow through was transferred to a fresh 1.5 mL tube and centrifuged for 20 minutes at room temperature and 5000 rpm. Pellets were resuspended in 100 uL 1x SDS Loading Buffer and 50 uL of 2M Tris pH 7.6 was added to neutralize the extract. Samples were boiled for 10 minutes, centrifuged at max speed for 10 minutes, and supernatant was used immediately to load onto SDS-PAGE gels prior to Western blotting or flash frozen and stored at -80°C.

### Yeast Chromatin Enriched Fractions (yCHEFs)

yCHEFs was performed as previously described with minor modifications.^121^ All samples were completed in biological replicate in isogenic strains derived from independent parental strains. 120 OD units of purified quiescent cells were pelleted and were resuspended in 200 uL Buffer 1 (20mM HEPES pH 7.6, 60 mM KCl, 15 mM NaCl, 10 mM MgCl_2_, 1 mM CaCl_2_, 10 mM N-butyric acid, 0.8% Triton X-100, 0.25 M Sucrose, 2.5 mM spermidine, 0.5M spermine, 1xPMSF) and transferred to tubes containing ∼200 uL acid washed glass beads. The cells were then lysed by bead beating. To collect cell lysates, a needle was used to make a hole in the bottom of the tube and tubes were placed in 15 mL conical tubes. Samples were centrifuged at 600 rpm for 2 minutes at 4°C. Lysates were transferred to 1.5 mL tubes and centrifuged at 500 rcf for 5 minutes at 4°C. A 20 uL aliquot of the supernatant was taken and placed on ice as the whole cell lysate fraction. The remaining supernatant was transferred to new 1.5 mL tubes and centrifuged at max speed for 20 minutes at 4°C. Pellets were resuspended in 200 µL Buffer 1 and centrifuged again at max speed for 20 minutes at 4°C. Pellets were then resuspended in 200 µL Buffer 2 (20 mM HEPES pH 7.6, 450 mM NaCl, 7.5 mM MgCl_2_, 20 mM EDTA pH 8, 10% Glycerol, 1% IGEPAL, 0.5 M sucrose, 2 M urea, 1 mM DTT, 1x PMSF) and centrifuged at max speed for 30 minutes at 4°C. Pellets were resuspended in preheated 30 µL 1x loading buffer without dye. For whole cell lysates, 20 µL preheated 2x SDS loading buffer without dye was added. Samples were boiled for 10 minutes and centrifuged at max speed for 2 minutes at 4°C. Supernatants were transferred to new tubes, flash frozen, and stored in -80°C. Protein concentrations were quantified using the Qubit Protein Assay kit and 46 µg of each sample was loaded onto 10% SDS-PAGE gels prior to Western blotting.

### Confocal microscopy and chromatin volume quantification

Quiescent cells from *HTB2-GFP* strains were purified as described above. All samples were performed in biological replicate in two isogenic strains derived from independent parent strains. A 1.7% agarose solution in water was used to prepare ∼1 cm^2^ square agarose pad on microscope slides. One µL of cell suspension was layered on top of the pad and spread out with a microscope coverslip. Full Z stacks (31 planes with 0.2 µm spacing) of GFP-labeled chromatin were acquired. Images were collected on a Nikon Ti-E microscope equipped with a 100X (NA: 1.49) TIRF objective, a Ti-S-E motorized stage, piezo Z control (Physik Instrumente), an iXon DU888 cooled EM-CCD camera (Andor) and a spinning disc confocal scanner unit (CSUX1, Yokogawa). A 488 nm laser housed in an LU-NV laser unit equipped with AOTF control (Nikon) was used to excite GFP, which was used with an emission filter mounted in a filter wheel (ET525/50M; Chroma). The microscope was controlled with NIS Elements (Nikon). The same settings were used for imaging all cells and all images were taken on the same day to ensure consistency. Nuclear volumes were measured using FIJI.^122^ One threshold that captures cells in all Z stacks was used and chromatin volumes were measured using 3D object counter. Objects less than 50 voxels were excluded from the analysis. Outliers lower than Q1−1.5 IQR or greater than Q3 +1.5 IQR were removed. Significance was determined using either one-way ANOVA followed by Bonferroni’s multiple comparisons test or unpaired t-test with Welch’s correction via GraphPad Prism.

### ChIP-seq

Chromatin immunoprecipitation was performed as described with modifications.^123^ All ChIP samples were performed in two biological replicates in isogenic strains derived from independent parental strains. 1000 OD units of purified quiescent cells or 300 OD units of cycling cells were crosslinked with formaldehyde at a final concentration of 1% for 1 hour at room temperature on an orbital shaker. Crosslinking was quenched for 5 minutes by addition of glycine to 0.125 M. Cells were washed with cold TBS, flash frozen, and stored at -80°C. For chromatin preparation, pellets were resuspended in cold Breaking Buffer (100 mM Tris pH 8, 20% glycerol) supplemented with protease inhibitor cocktail to 1x (100x protease inhibitor cocktail: 100 mM PMSF, 200 µM pepstatin, 60 µM leupeptin, 200 mM benzamidine, 200 µg/mL Chymostatin A) and lysed by bead beating with ∼250 µL acid washed glass beads for 6 minutes. Cell lysate was collected in low-retention 1.5 mL tubes and centrifuged at maximum speed for 1 minute. Pellets were resuspended in 1 mL FA Low Salt Buffer (50 mM HEPES-KOH pH 7.6, 150 mM NaCl, 1 mM EDTA, 1% Triton X-100, 0.1% sodium deoxycholate) supplemented with protease inhibitor cocktail. The suspension was transferred into 15 mL Diagenode tubes with sonication beads and sonicated using a Diagenode Bioruptor Pico sonicator for 23 cycles of 30 seconds on/30 seconds off. Samples were then centrifuged twice at 16,000 rcf for 10 minutes at 4°C, taking the supernatant each time. After the last centrifugation step, chromatin was flash frozen and stored at -80°C. To quantify the DNA, a 50 µL aliquot was taken from the chromatin preparation and added to 50 µL 2x Stop buffer (20 mM Tris pH 8.0, 100 mM NaCl, 20 mM EDTA, 1% SDS) and incubated at 65°C overnight. 2 µL of 10 mg/mL RNase A was added and samples were incubated for 1 hour at 42°C. Next, 2 µL of 10 mg/ml Proteinase K was added and samples were incubated for 4 hours at 55°C. DNA was purified using the Qiagen MinElute PCR Cleanup Kit (Qiagen, catalog #28004) and DNA was quantified via Qubit.

For immunoprecipitation (**IP**), 20 µL of Protein G Dynabeads (Invitrogen, catalog #10004D) per sample were washed three times in PBST and resuspended in 20 µL PBST per sample. Antibodies were conjugated to beads for 1 hour at 20°C at 1200 rpm. Beads were washed three times in PBST, one time in FA buffer, and resuspended in 20 µL of FA buffer per reaction. 1 µg of chromatin was combined with FA Buffer to a total volume of 350 µL and 46.67 µL was removed and reserved for input. 20 µL beads were added to the IP sample, and samples were incubated for 90 minutes with end over end rotation at room temperature. Beads were washed on a magnetic rack three times in Low-Salt FA buffer, three times in High-Salt FA buffer (50 mM HEPES-KOH pH 7.6, 500 mM NaCl, 1 mM EDTA, 1% Triton X-100, 0.1% sodium deoxycholate), and one time in RIPA buffer (10 mM Tris pH 8.0, 0.25 M LiCl, 0.5% IGEPAL, 0.5% sodium deoxycholate, 1 mM EDTA). DNA was eluted from beads twice by resuspension in 50 µL 2x stop buffer and incubation at 70°C for 15 minutes. 46.67 µL 2x Stop Buffer was added to reserved inputs, and IP and input samples were incubated at 65°C overnight to remove crosslinks. Samples were then treated with RNase A and Proteinase K and DNA was purified as described above. Libraries were prepared using the Tecan Ovation Ultralow V2 kit (NuGEN, catalog #0344). H3 IP samples were amplified for 14 cycles and all other IP samples were amplified for 16 cycles. Inputs were amplified for 13 cycles. Samples were sequenced using 150 bp paired-end sequencing via the NovaSEQ X platform with a target of 10 million reads per sample.

### ChIP-seq data analysis

Fastq files were aligned to the sacCer3 reference genome using bowtie2.^124^ Samtools was used to generate sort, and index bam files.^125^ deeptools2 was used to convert bam files to reads per kilobase per million (**RPKM**) normalized bigWig files using the bamCoverage command with the options --normalizeUsing RPKM --ignoreDuplicates.^126^ Normalization to input was done using deeptools2 bigwigCompare. ComputeMatrix reference-point was used to generate matrices for plotting using plotHeatmap or plotProfile. For each ChIP sample, data from each of two replicates were examined for agreement by eye and Pearson correlation, then replicate data were merged and reanalyzed to increase signal to noise in the final analysis. Peak calling was done on merged samples with MACS2 using the following command: macs2 callpeak -t IP_file.bam -c IN_file.bam -n out_file_name -f BAMPE -g 1.21e7 -B --nomodel --extsize 147.^62^ To find differential peaks (represented in bar graphs), the following command was used: macs2 bdgdiff --t1 AID treat_pileup.bdg --c1 AID control_lambda.bdg --t2 Control treat_pileup.bdg --c2 Control control_lambda.bdg --d1 AID depth --d2 control depth - g 60 -l 147 --o-prefix out_file_name_. To find overlaps between peaks (represented in Venn diagrams), the bedtools intersect command was used.^63^ Overlap analysis was conducted on peak center +/-1kb for Brn1 and Rpb1, and on peak center +/- 500bp for Brn1 and RPA.

### Micro-C XL

Micro-C XL was completed as described with modification.^127^ All samples were performed in biological replicate in isogenic strains derived from independent parental strains. 8000 OD units of purified quiescent cells were crosslinked with 3% formaldehyde in ultrapure water in a 1.2 L final volume for 10 min at room temperature. Crosslinking was quenched by addition of 130 mL of 2.5 M glycine. Cells were centrifuged at 3000 rpm at 4°C for 5 min and washed once with ultrapure water. For spheroplasting, cell pellets were resuspended in 120 ml Buffer Z (1 M sorbitol, 50 mM Tris pH 7.4, 10 mM β-mercaptoethanol) and the cell suspension was distributed to twelve 15 mL conical tubes. 1 mL of 10 mg/ml Zymolyase 100T (Amsbio, catalog #120493-1) was added to each tube and cells were incubated at 30°C with rotation until ∼80% of cells were spheroplasted (approximately 2 hours). Spheroplasts were centrifuged at 3000 rpm at 4°C for 10 min and washed gently without resuspension in cold PBS. For long-range crosslinking, DSG (ThermoFisher, catalog #PI20593) was dissolved in DMSO to obtain a 300 mM solution and subsequently diluted to 3 mM in PBS. Spheroplast pellets were resuspended in 5 mL 3mM DSG (ThermoFisher, catalog #PI20593) crosslinking solution and incubated at 30°C with rotation for 40 min. Crosslinking was quenched with 1 mL 2.5M glycine and spheroplasts were centrifuged at 3000 rpm for 10 minutes at 4°C. Without disrupting the pellet, spheroplasts were washed with cold PBS and the pellets were flash frozen and stored at -80°C. For each sample, Microccocal nuclease (**MNase**) (Worthington, catalog #LS004798) titrations were performed on spheroplasts from 2 pellets split into 8 MNase titration reactions. The MNase concentration yielding ∼90% mononucleosomes and 10% dinucleosomes was chosen to complete the rest of the Micro-C protocol. The remaining 10 pellets were used to perform the Micro-C protocol as described,^60^ completing the end repair, biotin labeling, and ligation reactions in 10 replicates, one per each tube, prior to combining samples into two for the dinucleosomal DNA purification and into one for the library prep. Libraries were generated using the NEBNext Ultra II DNA Library Prep Kit for Illumina (NEB, catalog #E7645) with modifications. 2.5 µL MyOne Streptavidin C1 beads (ThermoFisher, catalog #650.01) were washed with 1x TBW buffer (0.1% Tween20, 5 mM Tris-HCl pH 7.5, 0.5 mM EDTA, 1 M NaCl) at room temperature with rotation and resuspended in 150 µL 2x BW buffer (10 mM Tris pH 7.5, 1 mM EDTA, 2 M NaCl) and used to purify biotinylated DNA. 150ul of the biotinylated DNA diluted in 0.1X TE was added to beads (final volume 300 µl) and incubated at room temperature with rotation for 20 min. Beads were washed once with TBW buffer (0.1% Tween20, 5 mM Tris-HCl pH 7.5, 0.5 mM EDTA, 1 M NaCl) at room temperature with rotation, once with TBW at 55°C with 1000 RPM shaking, and once with NEBuffer 2 at room temperature with rotation. Beads were resuspended in 47 µL 1x TE, and on-bead end repair and ligation reactions were done according to manufacturer protocol. For DNA clean up, beads were washed once with TBW buffer at room temperature with rotation, once with TBW at 55°C with 1000 RPM shaking, and once with 0.1x TE at room temperature with rotation. Beads were resuspended in 16 µL 0.1x TE buffer. Before amplifying the library, a test PCR was performed on 1 µL of bead solution for 11 cycles. PCR product was purified and size selected using Cytiva SeraMag Select beads (Cytiva, catalog #29343045) added at a 0.9x beads to sample ratio. The DNA concentration was measured via Qubit 1x HS dsDNA Assay and the number of cycles to amplify the remaining DNA was calculated by determining the least number of cycles expected to yield 80 ng total DNA. Samples were sequenced using the Illumina NovaSeq X platform, obtaining ∼300 million 150 bp paired-end reads per replicate.

### Micro-C XL data analysis

Biological replicates were analyzed, found to agree by HiCRep,^88^ then merged and reanalyzed to increase signal to noise in the final analysis. Fastq files were trimmed to 50 bp using fastp.^128^ Paired fastq files were aligned to the sacCer3 reference genome using bwa-mem2.^129^ Reads with MAPQ score < 2 were discarded. Sam files were converted to BAM files using samtools view.^125^ Ligations were detected and analyzed using the Pairtools pipeline.^130^ Pairtools parse was used to detect ligation junctions and generate pairs files, followed by pairtools sort and pairtools dedup for duplicate removal. Uniquely mapped valid interactions were retained using pairtools select. Uniquely ligated reads were separated based on orientation (IN-IN, IN-OUT, OUT-IN, and OUT-OUT). IN-IN reads were excluded from all further analysis to ensure all pairs resulted from MNase digestion and ligation. Micro-C contact maps were generated using juicer and visualized with Juicebox.^131^ The Cooler package was used for iterative correction (**ICE**) balancing of contact matrices and to generate multiple resolution mcool files.^132^ Stratum adjusted correlation coefficient SCC scores were calculated with contacts within 90 kb to assess reproducibility using the HiCRep package via the command line.^88^

The probability of contact P(s) relative to genomic distance was determined using cooltools with the command: cooltools expected-cis $file.cool –view $view_file.tsv -o $New_file.tsv –smooth –aggregates-smoothed –smooth-sigma 0.1. Contact probability from 300 bp to 2 Mb was included in the analysis. L-CID boundaries and insulation scores were called using the cooltools command: cooltools insulation –threshold 0.4 -o $out-file.tsv –view $view_file.tsv $input-file.cool. Boundary overlaps were determined with the following bedtools command: bedtools intersect -a $boundary-file1.bed -b $boundary-file2.bed -u > $boundaries-in-both.bed. Boundary overlaps with ChIP-seq peaks were determined using bedtools with the following command: bedtools window -a $boundary-file.bed -b $condenisn-peak-file.bed -w 1000 -u > $boundaries-overlapping-condensin.bed.

Pileups centered on Brn1 peaks were generated using the coolpuppy package with the following command: coolpuppy.py $file.cool $condensin_peaks.bed -o $out-file.clpy –pad 10000.^133^ 600 resolution pileups were generated using: coolpuppy.py $file.mcoo::/resolutions/600 $condensin_peaks.bed -o $out-file.clpy --pad 10000. Division pileups were generated using: dividepups.py $file1.clpy $file2.clpy -o $division-file.clpy. Local pileups were generated using: coolpuppy.py $file.cool $condensin_peaks.bed -o $out-file.clpy – LOCAL –pad 30000. Loop piles were generated using: coolpuppy.py $file.cool $condensin_loops.bedpe -o $out-file.clpy –pad 10000.

---

**Figure S1.**
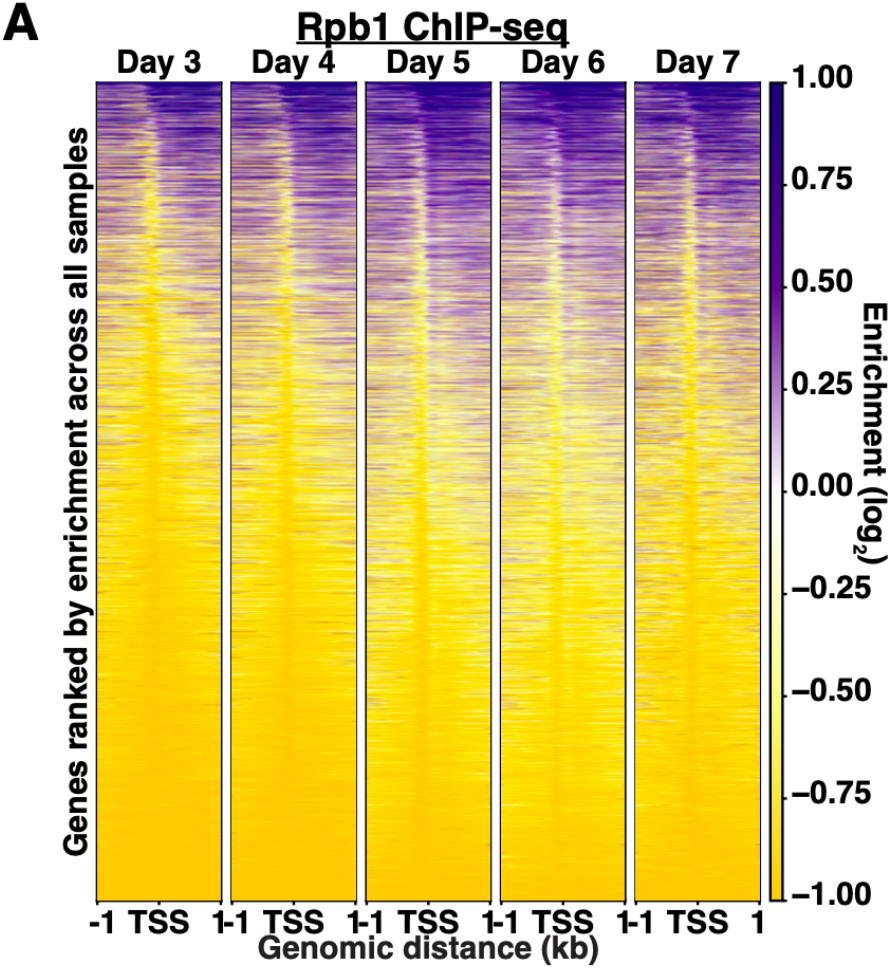
Rpb1 occupancy is dynamic during quiescence entry. (**A**) ChIP-seq heatmaps of Rpb1 in quiescent cells purified on the indicated day.

**Figure S2.**
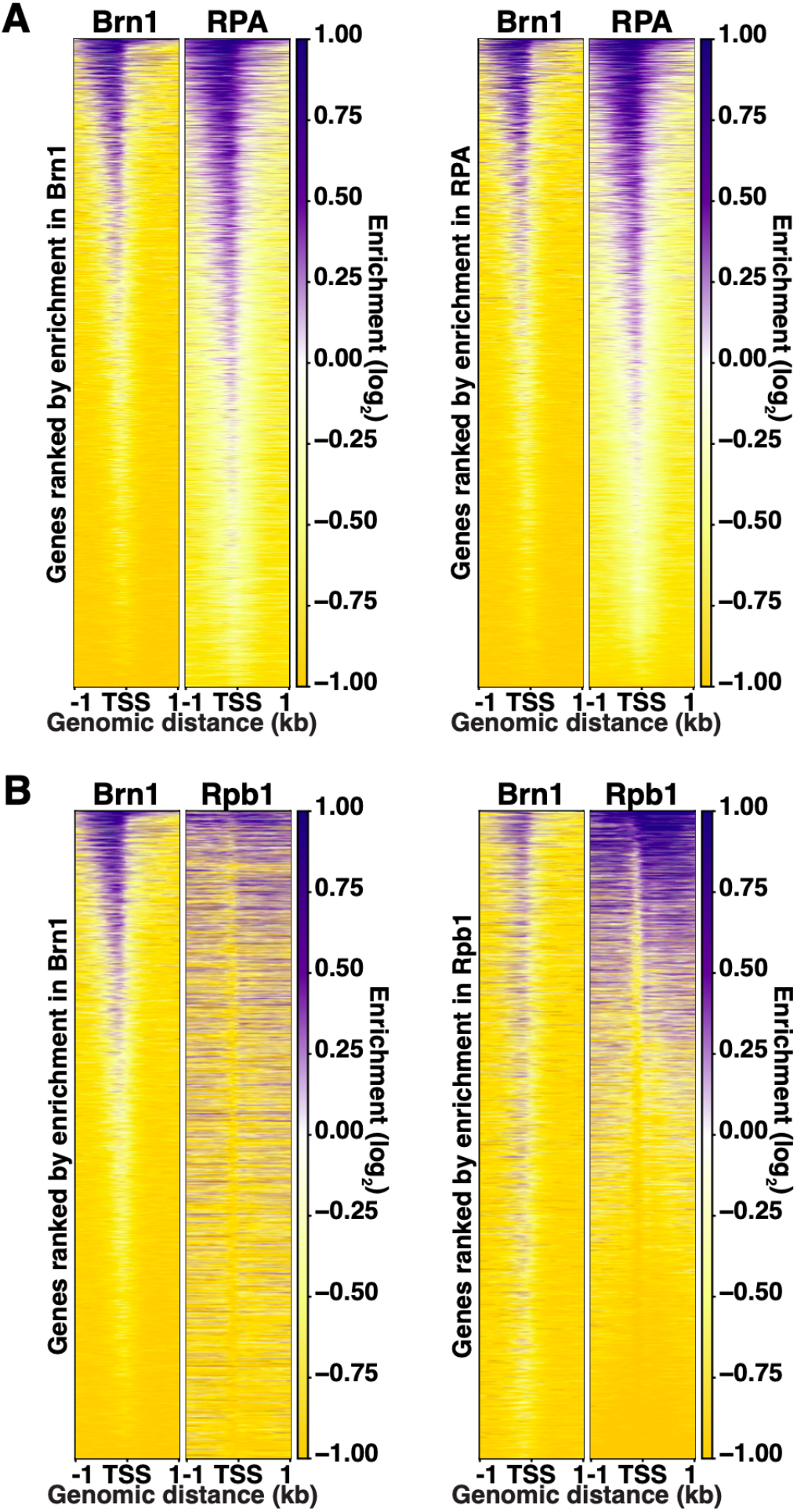
Condensin localization correlates with RPA more than RNAPII. (**A**) ChIP-seq heatmaps of Brn1-FLAG and RPA in Day 7 cells sorted by either Brn1 or RPA occupancy. (**B**) ChIP-seq heatmaps of Brn1-FLAG and Rpb1 in Day 7 cells sorted by either Brn1 or Rpb1 occupancy.

**Figure S3.**
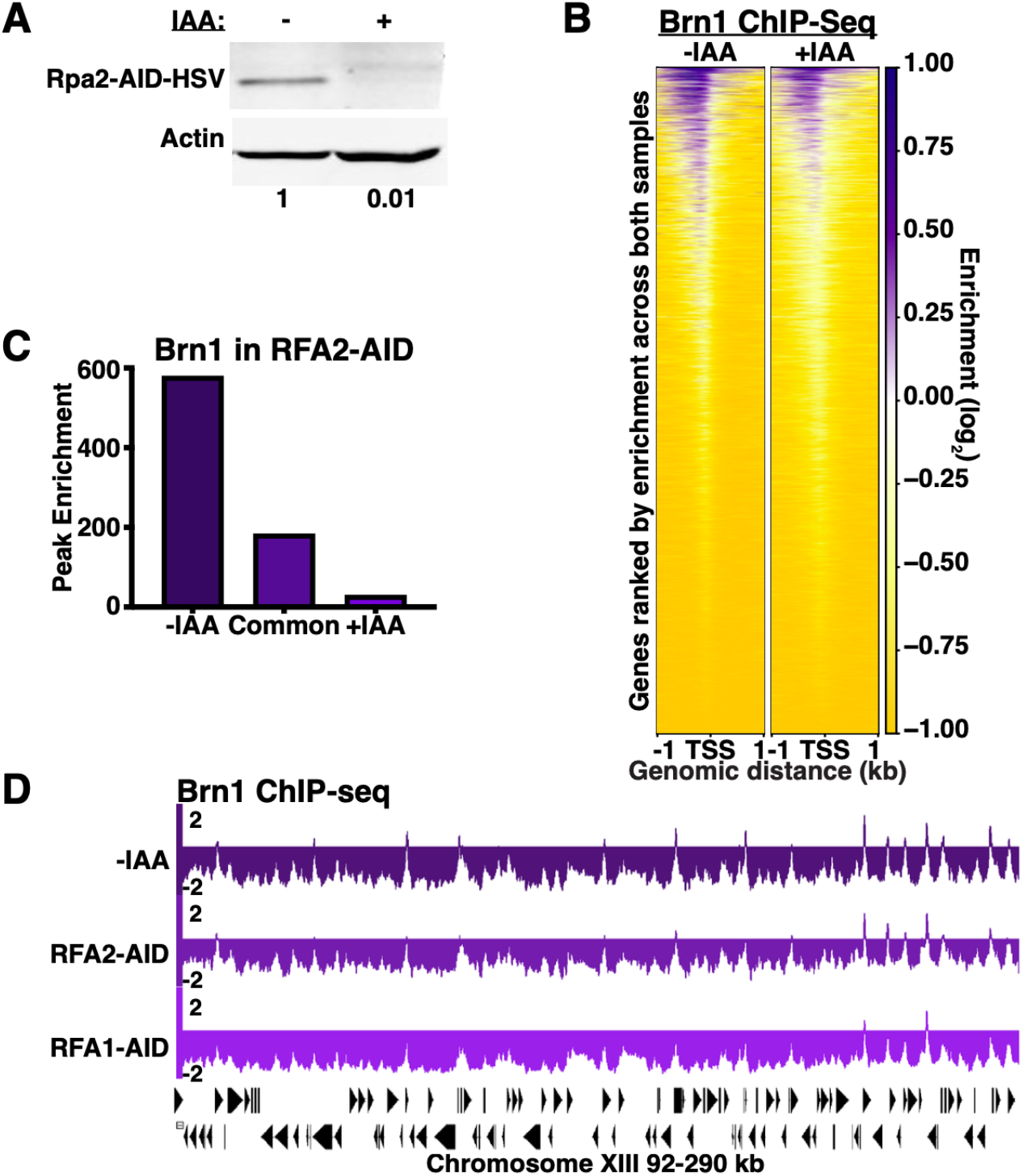
The Rpa2 subunit of RPA is required for condensin targeting during quiescence. (**A**) Representative Western blot in *RFA2*-AID-HSV Day 7 cells +/-IAA. Band is normalized to actin and then to -IAA. (**B**) ChIP-seq heatmaps of Brn1-FLAG in *RFA2*-AID-HSV Day 7 cells +/- IAA. (**C**) Differential peak analysis of Brn1-FLAG in *RFA2*-AID-HSV Day 7 cells +/- IAA. (**D**) Representative genome browser view of ChIP-seq data.

**Figure S4.**
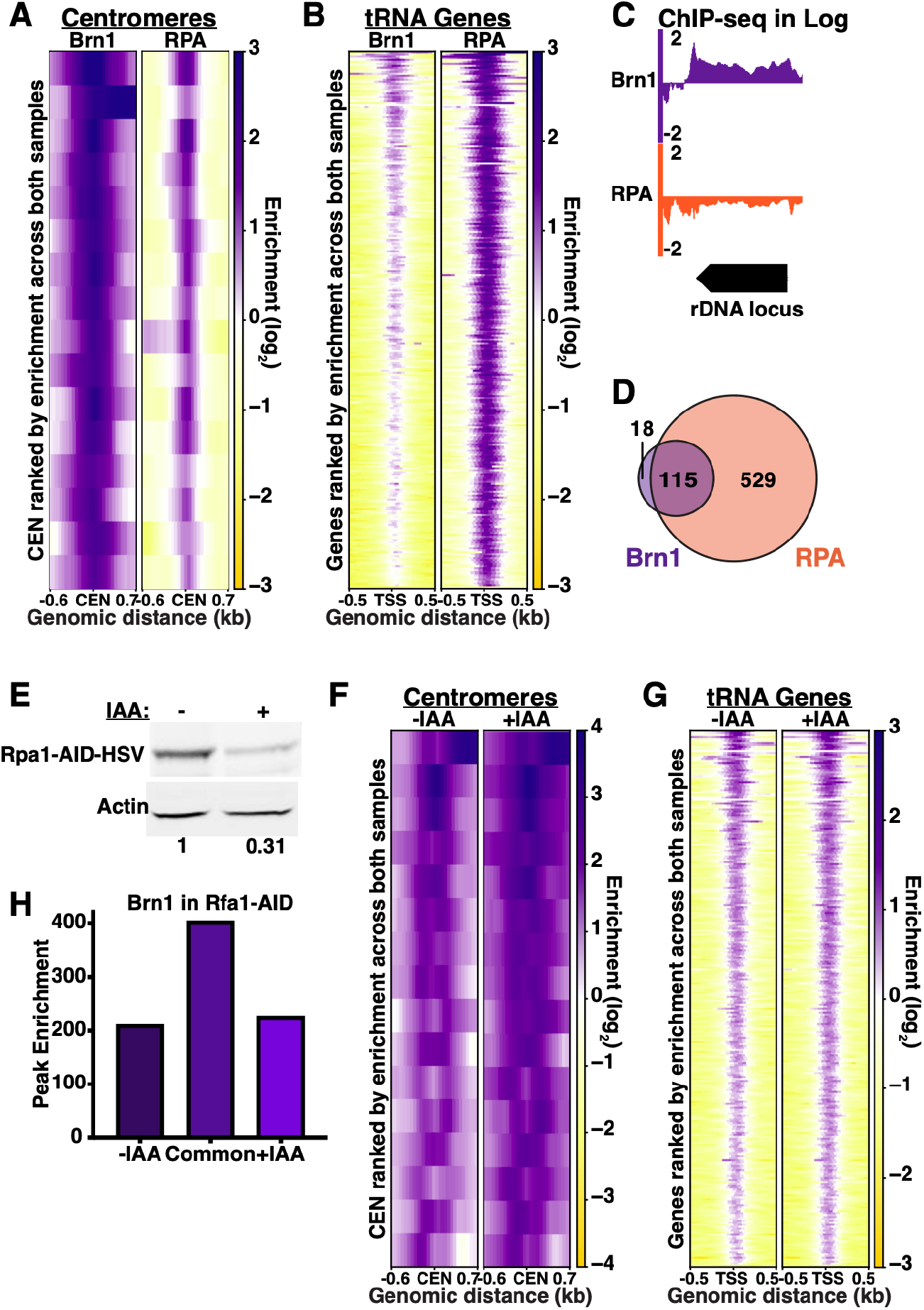
RPA does not target condensin in cycling cells. (**A**) ChIP-seq heatmaps of Brn1 and RPA at centromeres in cycling (**log**) cells. (**B**) ChIP-seq heatmaps of Brn1-FLAG and RPA at tRNA genes in log cells. (**C**) Genome browser view of Brn1-FLAG and RPA at the rDNA in log cells. Occupancy over rDNA repeats is averaged and represented as a single locus. (**D**) Overlap of ChIP-seq peaks filtered using a 1.5-fold enrichment threshold. Peaks within 1 kb were determined to overlap. (**E**) Representative Western blot in *RFA1*-AID-HSV log cells. Band was normalized to actin and then to -IAA. (**F**) ChIP-seq heatmaps of Brn1-FLAG at centromeres in *RFA1*-AID-HSV log cells +/-IAA. (**G**) ChIP-seq heatmaps of Brn1-FLAG at tRNA genes in *RFA1*-AID-HSV log cells +/-IAA. (**H**) Differential peak Brn1-FLAG in *RFA1*-AID-HSV log cells +/-IAA.

**Figure S5.**
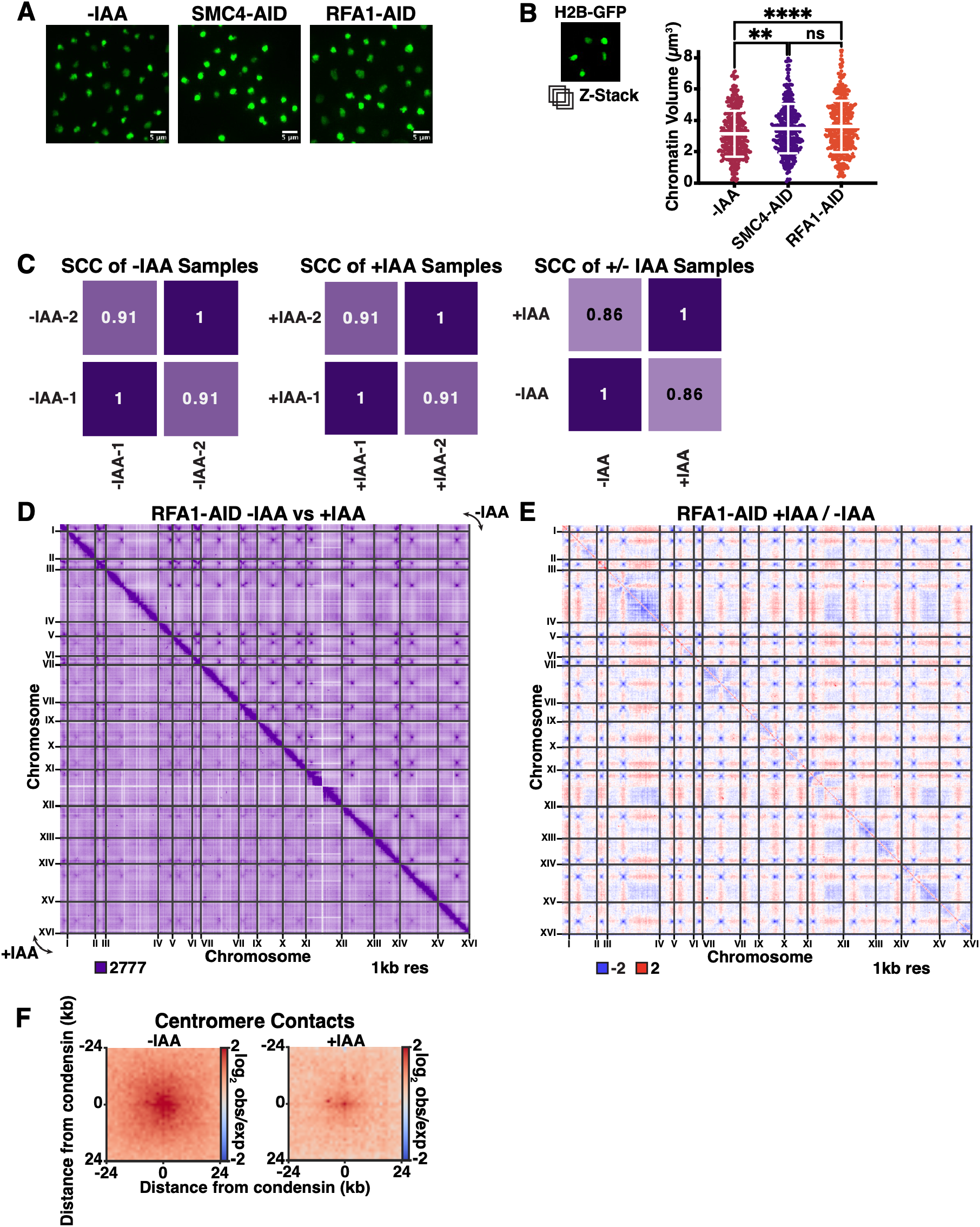
Loss of RPA leads to large-scale chromatin reorganization. (**A**) Representative confocal microscopy images of Day 7 cells +/-IAA in the indicated strains. (**B**) Chromatin volume quantification of Z-stacks. Bars represent mean +/- SD. Each time point consists of at least 150 cells from each of two biological replicates. Significance measured using one-way ANOVA. (**C**) SCC scores of Micro-C replicates and merged samples calculated using HiCRep. (**D**) Genome-wide Micro-C contact map in *RFA1*-AID Day 7 cells +/- IAA. (**E**) Genome-wide Micro-C contact map in *RFA1*-AID Day 7 cells. +IAA data are divided by -IAA data. (**F**) Pileup plots of Micro-C contacts between centromere pairs in *RFA1*-AID Day 7 cells +/- IAA.

**Figure S6.**
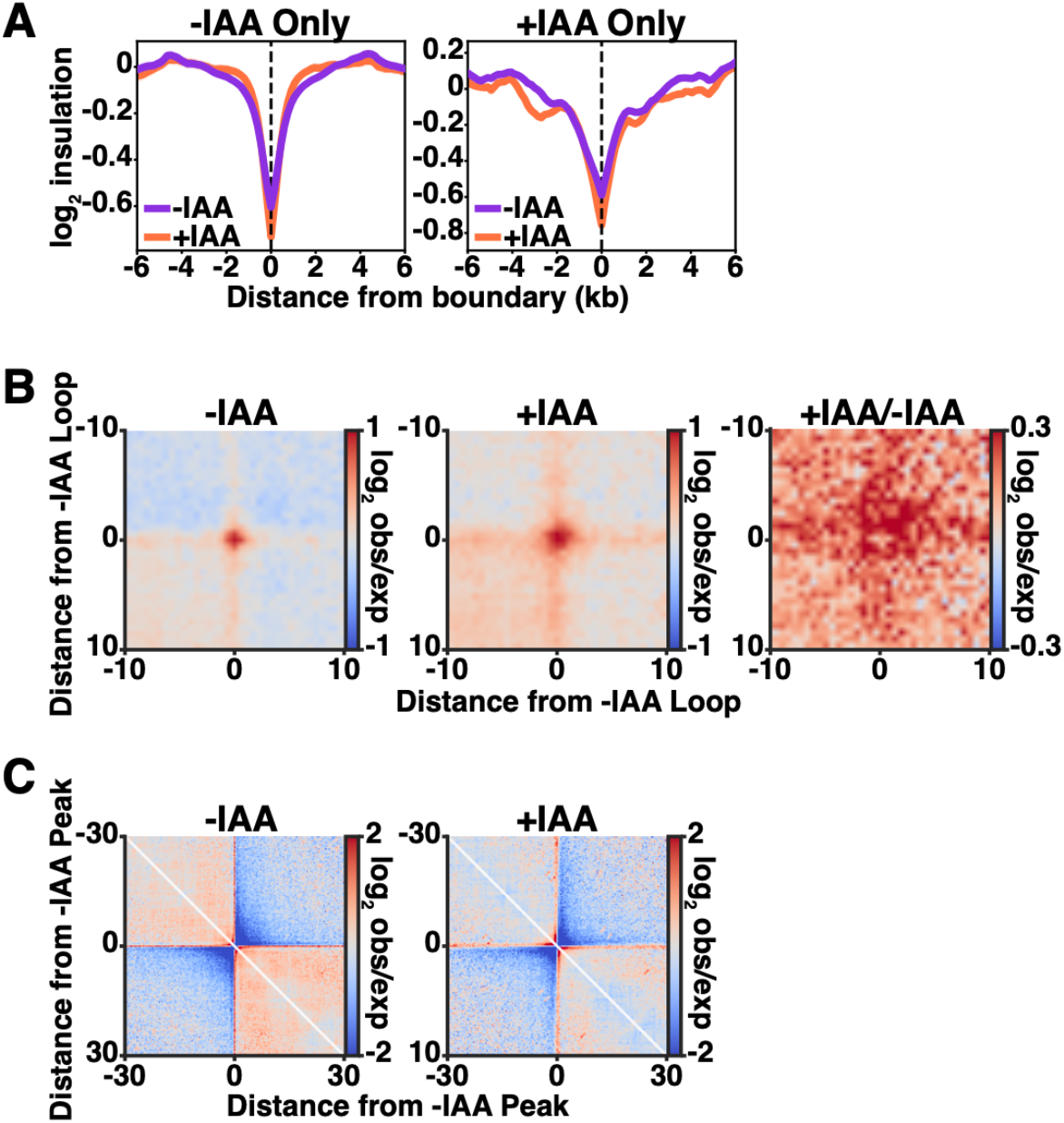
RPA loss leads to alterations in condensin-mediated 3D chromatin structure. (**A**) Insulation plots centered at L-CID boundaries overlapping Brn1 peaks in either -IAA or +IAA. (**B**) Pileup plots of 150 bp Micro-C data at loops called in RFA1-AID Day 7 cells -IAA. Right plots show +IAA data divided by -IAA data. (**C**) Local pileup plots of 150 bp Micro-C data centered on Brn1 peaks called in *RFA1*-AID Day 7 cells -IAA.

